# A hierarchical nickel organic framework confers high conductivity over long distances in cable bacteria

**DOI:** 10.1101/2025.10.10.681601

**Authors:** Filip J. R. Meysman, Bent Smets, Silvia Hidalgo Martinez, Nathalie Claes, Bob C. Schroeder, Jeanine S. Geelhoed, Yun Liu, Jiji Alingapoyil Choyikutty, Tamazouzt Chennit, Thijs Bodson, Alberto Collauto, Maxie M. Roessler, Dmitry Karpov, Sylvain Bohic, Matteo Aramini, Shusaku Hayama, Maxwell Wetherington, Martijn A. Zwijnenburg, Galina Pankratova, Isabel Pintelon, Jean-Pierre Timmermans, Gert Nuyts, Karolien De Wael, Sara Bals, Jo Verbeeck, Han Remaut, Henricus T. S. Boschker

## Abstract

Multi-cellular cable bacteria have evolved a unique machinery that efficiently transports electrons across centimetre-scale distances. Currents flow through a parallel network of periplasmic fibres, which display an extraordinary conductivity for a biological material. However, the conduction mechanism remains elusive as the molecular structure of the fibres has not been resolved. Here, we demonstrate that each fibre embeds a bundle of intertwined nanoribbons, which are built from Nickel Bis(Dithiolene) (NiBiD) repeat units that are formed by linking nickel centres with ethenetetrathiolate ligands. The planar and conjugated NiBiD complexes are aligned and stacked to form an elongated supramolecular coordination network, thus explaining the observed organo-metal electronic properties of the fibres. Our results hence demonstrate that biology is capable of producing extensive metal organic frameworks. These structures enable highly conductive one-dimensional conduits, ensuring efficient charge transport over macroscale distances, thus providing a novel design principle for bio-based, sustainable organo-electronic materials.

## Main text

Cable bacteria stand out in the prokaryotic world for their unique capability to channel electrical currents over distances spanning several centimeters^1–4^. To enable such long-range conduction, these filamentous bacteria contain a regularly spaced network of fibres embedded in the cell envelope^5^. These periplasmic fibres run continuously along the entire centimetre-long filament and exhibit remarkable electrical properties, including an electrical conductivity (> 100 S/cm) that exceeds that of highly doped, synthetic semiconductors^6–9^. However, the mechanism by which these fibres attain such extraordinary conductance remains unresolved.

Electron transfer through proteins generally requires the presence of non-protein cofactors that act as relay centres for electrons to pass through the otherwise insulating protein matrix. These juxtaposed cofactors are known to facilitate electron transport from the nanometre scale, as seen in the membrane complexes involved in photosynthesis and respiration^10,11^, up to the micrometre scale, as encountered in the multi-heme cytochrome-containing appendages of metal reducing bacteria^12–13^. The multi-step hopping of electrons through such protein systems generally relies on Fe-containing cofactors, such as Fe-S clusters or hemes^11,14^.

Conduction in cable bacteria, however, spans a much larger distance and does not seem to conform to an Fe-based conduction mechanism. Raman spectroscopy indicates that the periplasmic fibres cannot contain Fe-S clusters nor hemes^6^. Instead, distinct vibrational modes indicate the presence of a metallocentre containing nickel, while ^34^S isotope labelling experiments show that the Ni centre is sulfur-ligated^15,16^. The proposition of a Ni/S-dependent conduction mechanism is remarkable as Ni is known to be included in only nine enzymes, and in each of these, the Ni-site serves a catalytic function and not an electron relay function^17^.

Hitherto, the molecular structure of the Ni/S complex remains unresolved. Here, we applied detailed microscopy and spectroscopy, combined with Density Functional Theory (DFT) calculations, to establish a molecular basis for the efficient, long-range electron transport in cable bacteria. Our results demonstrate that the Ni/S entity comprises an oligomeric nickel bis(1,2-dithiolene) complex, of which the repat units are assembled into a one-dimensional, metal-organic framework, thus producing nanoribbons that run inside the conductive fibres. Such metal-organic frameworks are not known to be biologically produced, but synthetic versions are attracting considerable interest within the field of organic electronics^18^, as they show exceptional opto-electronic properties^19–21^. Consequently, it appears that already long ago, microbial evolution has explored the potential of metal-organic frameworks to attain extended, flexible structures with high conductivity.

### Fibres contain an intertwined bundle of Ni-rich nanoribbons

To document the substructure of the internal fibre network, cable bacterium filaments were individually isolated from sediment incubations. Scanning Electron Microscopy (SEM) shows that the outer surface of the filaments displays regularly spaced, parallel ridges (width 134 ± 3 nm; n=248), while local constrictions indicate the presence of internal junctions between cells (Fig. 1a,b). Transmission Electron Microscopy (TEM) of resin-embedded cross-sections (Fig. 1c) reveals that each ridge harbours a periplasmic fibre (diameter 49 ± 5 nm; n=50), which appears as an unstained patch, thus suggesting a dense structure resistant to diffusional penetration of the stain (Fig. 1d; N_F_=68 fibres per filament).

**Fig. 1:**
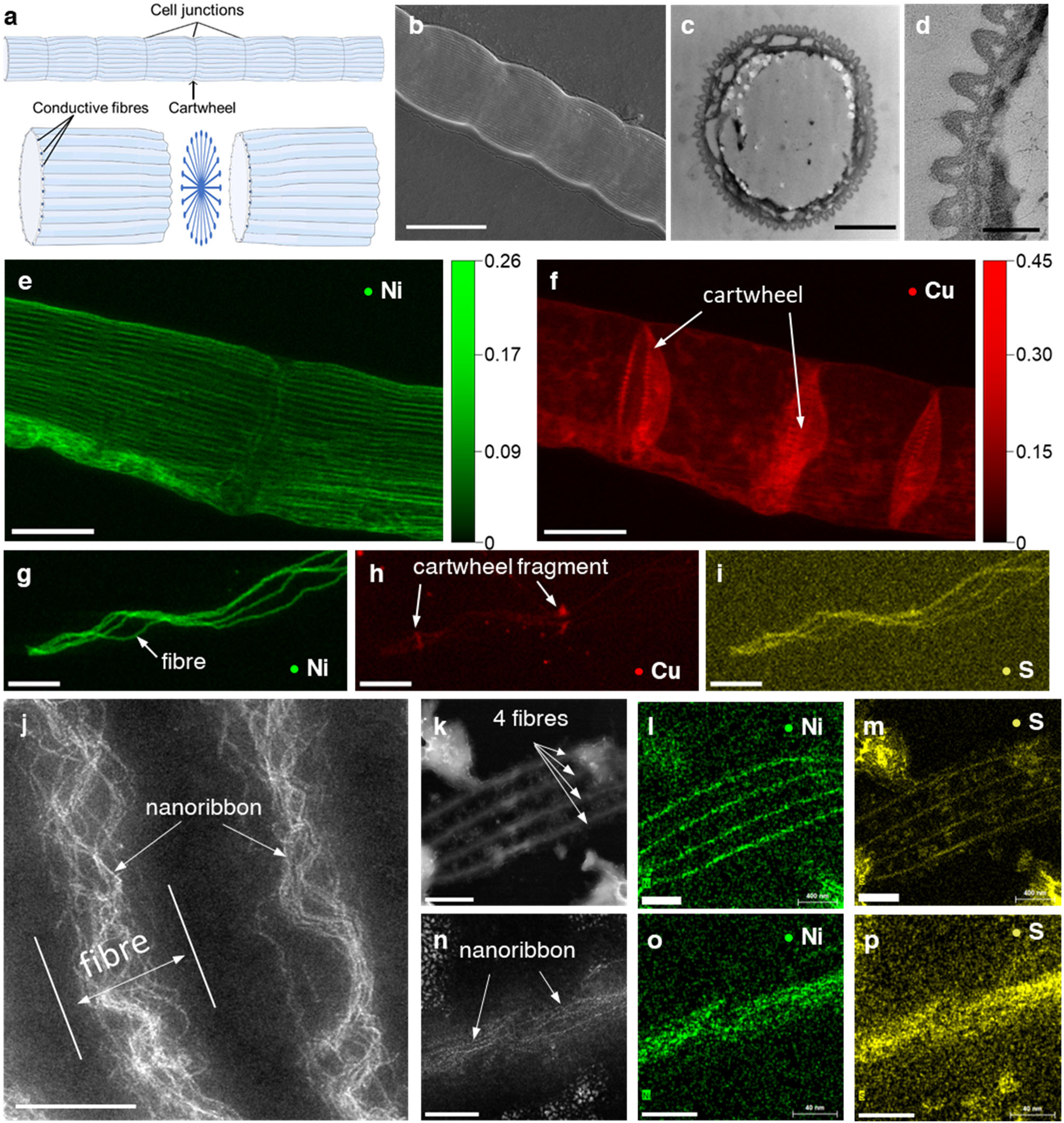
Microscopic imaging and nXRF mapping of periplasmic fibres in cable bacteria. **a**, Schematic of filament morphology. **b**, SEM image of a native filament (HV: 20 kV, WD: 10 mm, Mag: 7960x) showing parallel ridge compartments. Scale bar: 4 µm. **c**, TEM image of cross-section of a resin-embedded native filament. Scale bar: 1 µm. **d**, Detail of panel c showing single fibres (white patch) embedded within ridge compartments. Scale bar: 200 nm. **e,f,** nXRF area density maps (ng mm^-2^) of Ni and Cu in a fibre skeleton. Scale bar: 2 µm. **g-i**, nXRF mapping of individual fibres. Elements are indicated. Scale bar: 2 µm. **j**, HAADF-STEM intensity image of a fibre skeleton piece with 2 parallel fibres revealing wavy, electron-dense channels (nanoribbons). Scale bar: 100 nm. **k-m**, HAADF-STEM-EDX images of a piece of fibre skeleton with 4 fibres. Scale bars: (K) 500 nm, (L) 400 nm (M) 400 nm. **n-p**, HAADF-STEM-EDX of a single fibre showing nanoribbons. Scale bars: 50 nm.

Sequential extraction of native filaments removes the lipid membranes and cytoplasm, thus producing a “fibre skeleton” ^6,15^ that retains the conductive fibre network implanted onto a connective sheath (Extended Data Fig. 1). We verified that fibre skeletons were equally conductive as native filaments^6^, thus demonstrating that the extraction does not notably affect the structure and transport properties of the fibre network. Nano-X-Ray Fluorescence (nXRF) imaging of native filaments (Extended Data Fig. 2a) and fibre skeletons (Fig. 1e; Extended Data Fig. 2b) reveals a distinct pattern of parallel lines in the Ni area density (ρA_Ni_) map, thus confirming the presence of Ni^15,16,22^. Lines are equidistantly spaced and remain continuous across the interfaces between cells. The regular spacing of the ρA_Ni_ peaks (147 ± 9 nm; n = 182) matches the geometry of the ridges as seen in SEM (Fig. 1b), and the full width at half maximum of the ρA_Ni_ peaks (52 ± 15 nm; n = 128) concurs with the fibre diameter as seen in TEM (Fig. 1c; Supplementary Fig. 1a).

The cell-cell interfaces are not enriched in Ni, but show an elevated Cu signal (Fig. 1f; Extended Data Fig. 2e,f) that traces the shape of the so-called “cartwheel” structure, which consists of a set of rod-like structures (“spokes”) that extend inwards from underneath each ridge to converge in a central node^5,22^. This cartwheel is also conductive, thus providing an electric interconnection between fibers and imposing redundancy on the electrical network^23^. Superposition of the Cu and Ni maps shows that the terminal ends of the Cu-rich spokes are connecting to the Ni-rich fibres (Supplementary Fig. 1b). There is no marked enrichment of iron (Fe) in the periplasmic fibres nor the cartwheel (Extended Data Fig. 2h), thus confirming that neither hemes nor Fe-S clusters are present^6,15^. When fibre skeletons were sonicated, loosened fibres and cartwheel fragments were produced, which reveal that sulphur (S) colocalizes with Ni in the fibres, and also with Cu in the cartwheels (Fig. 1g-i; Extended Data Fig. 2c,d; Supplementary Fig. 2-4). Our data hence suggest that the internal conductive network of cable bacteria incorporates two distinct metal complexes: a Ni/S-entity in the periplasmic fibres and a Cu/S-entity in the cartwheel spokes.

High-angle annular dark-field imaging (HAADF) scanning transmission electron microscopy (STEM) shows that each fibre harbours a set of electron-dense “nanoribbons” that follow a wavy trajectory along the longitudinal axis of the fibres (Fig. 1j; Supplementary Fig. 5). Energy-dispersive X-ray spectroscopy (EDX) confirms that these wavy ribbons are strongly enriched in Ni and S (Fig. 1k-p), thus suggesting a linear alignment of Ni/S complexes within a nanoribbon. Multiple ribbons (10-20) converge and diverge within each fibre, thus forming an intertwined channel network that flows in the direction of the fibres (Fig. 1j). Individual ribbons are 1.35 ± 0.25 nm wide (Supplementary Fig. 5d), and are confined within a central zone (∼30 nm wide) of the fibres. This aligns with previous Scanning Tunnelling Microscopy data, which suggested that the fibres contain a conductive core^15^. Detailed image analysis of nXRF and EDX maps enables quantification of Ni and S concentrations in the fibres and nanoribbons, accounting for S-containing amino acids in the surrounding protein structure (see S.I. for details; Supplementary Fig. 3-7). This constrains the molar S/Ni ratio of the wavy, electron-dense nanoribbons to range between 3.0 and 4.6 (Supplementary Tables 1-3).

### The Ni/S complex is a conjugated nickel bis(dithiolene)

Previously, we observed that the Ni/S-entity has a characteristic Raman fingerprint that is markedly different from known Ni-containing enzymes in biology, but shares many features with the Raman spectra of nickel bis(1,2-dithiolene) (NBDT) complexes^16^. These organic coordination compounds consist of a central nickel, coordinated by two bidentate 1,2-dithiolene ligands, thus forming a planar Ni(S_2_C_2_R_2_)_2_ complex with a D_2h_ symmetry and a delocalized electronic structure^24^. The R-substituents on the dithiolene ligands define the structure and properties of the NBDT complex (Extended Data Fig. 3a), but remained hitherto unresolved for the Ni/S-complex in cable bacteria^16^.

To confirm the NBDT-like character of the Ni/S complex, we compared the Raman spectra of fibre skeletons to an extensive set of synthetic NBDT reference compounds, which included six monomeric (one Ni centre) and two polymeric (multiple Ni centres) compounds (formulas in Extended Data Table 1; structures in Extended Data Fig. 3). While the Raman spectra of the Ni/S complex show marked differences with those of the monomeric NBDT (Extended Data Fig. 4), there was a tight resemblance to the spectra of the neutral coordination polymers poly-nickel(ethene tetrathiolate) (pNi(ett)) and poly-nickel(TetraThiaFulvalene tetrathiolate) (pNi(TTFtt)). The close correspondence between Raman peak positions (low-frequency peaks near 366 cm^-1^ and 494 cm^-1^ and thiocarbonyl radical (C=S⸱) peaks within 1150-1200 cm^-1^ region; Fig. 2a; Supplementary Table 5) and the similar wavelength dependence of the peak intensities (Supplementary Table 6) indicate that the Ni/S complex must share a high degree of structural homology with polymeric forms pNi(ett) and pNi(TTFtt) (see SI for additional discussion). Overall, the Raman data indicate that the Ni/S complex has an NBDT configuration with a limited number of bonds (Ni-S, S-C, C=C), and cannot feature nitrile groups or aromatic rings as R-substituents^25^. Moreover, the strong NIR resonance observed (Fig. 2a; Extended Data Fig. 4a) suggests that the complex is not monomeric^26^, but oligomeric or polymeric, in which multiple Ni centres are interconnected by dithiolene ligands to form a larger conjugated complex. We refer to this entity as the NiBiD (NiBisDithiolene) complex.

**Fig. 2:**
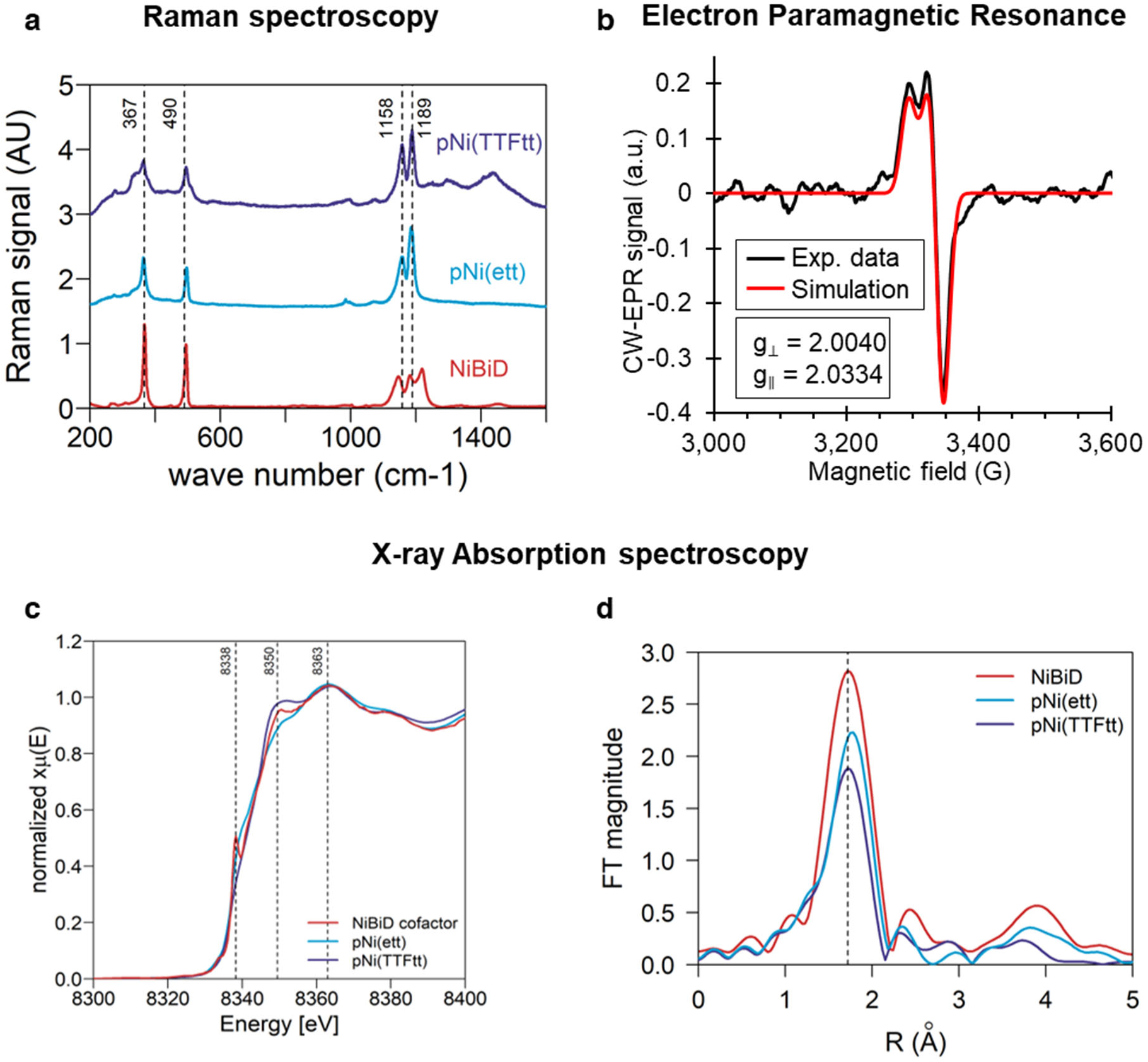
Spectroscopic investigation of the NiBiD complex. **a,** Raman spectra collected with 760 nm laser for two polymeric NBDT reference compounds pNi(ett) and pNi(TTFtt)) and cable bacterium fibre skeletons containing the NiBiD complex. a.u. = arbitrary units. **b**, Experimental and simulated EPR spectrum of native cable bacterium filaments at 12 K. **c**, The Ni K-edge XANES spectrum of native cable bacterium filaments containing the NiBiD complex compared to two polymeric NBDT reference compounds. **d**, Fourier transform magnitudes of the Ni K-edge EXAFS of native cable bacterium filaments compared to two polymeric NBDT reference compounds.

### The coordination environment of the Ni centre

The presence of thiocarbonyl radicals suggests a charge-neutral complex where dithiolene ligands are stabilized by ligand-centred radicals^20^. This hypothesis is supported by Electron Paramagnetic Resonance (EPR) spectroscopy of native cable bacterium filaments, which show a signal emerging at *g* ∼ 2 that is suitably simulated by an isolated S = ½ spin system (Fig. 2b). The average *g* value, *g*_av_ = (2·g_⊥_+g_∥_)/3 = 2.014, is consistent with an unpaired electron residing on an organic ligand rather than a transition metal, as also seen previously in pNi(ett) and pNi(TTFtt)^27^. The significant shift from the *g-*value of the free electron (*g*_e_ = 2.0023) indicates that the radical is centred on a heavier element like sulphur, which results in a larger spin-orbit coupling compared to lighter elements such as carbon, oxygen, or nitrogen. The low-field ‘shoulder’ can be accounted for by introducing a small axial anisotropy of the electron Zeeman interaction (Fig. 2b). The signal increases with decreasing temperature (Extended Data Fig. 5e) as expected for a centre with a paramagnetic ground state.

Spin quantification reveals spin concentrations less than one spin per Ni centre (0.12 ± 0.03, n=3; Extended Data Fig. 5c,d; Extended Data Table 2), aligning with values observed for poly-metal-(ett) structures^28^. DFT calculations show that when Ni(ett)-trimers are terminally capped with thiol groups or S – S bonded moieties (e.g., as crosslinked to cysteine residues) (structures III and IV in Extended Data Fig. 6), these structures are predicted to have an open-shell singlet ground state with two spins per oligomer (Supplementary Table 8). The open-shell character originates from two unpaired electrons that reside on the terminal ligands and is consistent with a S = ½ spin centre as suggested by the EPR signal. Congruent with the EPR data, there is negligible spin density on the nickel atoms, and the spins are predominantly localized on the sulphur and carbon atoms of the two external dithiolene ligands (Extended Data Fig. 6e). Conjointly, the spin data and DFT calculations support a NiBiD complex with multiple Ni centres (see SI Discussion).

To further constrain the coordination environment of the Ni centre, we applied X-ray Absorption Spectroscopy (XAS) to cable bacterium filaments under cryogenic conditions (10 K). As reference materials, we used inorganic Ni compounds with different oxidation states (Ni foil, pellets of NiO_2_ and NiO), the same NBDT reference compounds as investigated with Raman, and the Ni-pincer containing enzyme LarA^29^. The Ni K-edge of NiBiD lies at a different energy than the inorganic Ni reference compounds but closely resembles that of the NBDT complexes (Extended Data Fig. 7b). Consistent with a square planar coordination, NiBiD shows a pre-edge peak at 8,333 eV, which corresponds to the dipole-forbidden transition of the Ni 1s electron to the unoccupied 3d orbitals^30^ and a prominent rising-edge shoulder at 8,338 eV, which arises from a dipole-allowed 1s→4p_z_ transition^31,32^. NiBiD structurally differs from the Ni pincer in LarA^29^, as the spectrum of the latter shows a more pronounced pre-edge feature, a lower rising-edge shoulder, a more intense white line at 8,350 eV, and also markedly differs within the EXAFS region > 8,350 eV (Extended Data Fig. 7c). Among the NBDT compounds investigated, the XAS spectrum of NiBiD is most similar to that of the polymeric forms pNi(ett) and pNi(TTFtt), both in the XANES (Fig. 2c) and EXAFS (Fig. 2d) regions (additional discussion in SI). The same similarity was observed in the Raman spectra (Fig. 2a), thus suggesting a high level of structural homology.

### Axially aligned NiBiD complexes form 1D conduction channels

Combined, the Raman, EPR and XAS data support the hypothesis that NiBiD comprises an oligomeric form of either Ni_n_(ett)_n+1_ or Ni_n_(TTFtt)_n+1_, where n denotes the number of Ni centres. Both pNi(ett) and pNi(TTFtt) exclusively feature C-S bonds, and do not contain nitrile or benzyl groups, which hence must also be the case for NiBiD. The observed molar S/Ni ratio of 3.0-4.6 as determined by nXRF and STEM-HAADF-EDX puts an important constraint on the molecular configuration, and is only consistent with an ett-like structure (S/Ni ∼4) but not a TTFtt-like structure (S/Ni ∼8). A Ni_n_(ett)_n+1_ structure also adequately explains the fragmentation patterns previously obtained from ToF-SIMS analysis of fibre skeletons, which produces two specific organosulfur fragments (C_2_S_2_^−^ and C_2_S_2_H^−^) tightly associated with [NiS]_n_^-^ fragments^15^.

This invokes the question about the relative orientation and packing of NiBiD units within the electron-dense nanoribbons. The mean distance between Ni centres in Ni_n_(ett)_n+1_ is 0.603 nm (as derived from DFT model of a NiBiD unit; Extended Data Fig. 6), and so the observed Ni concentration in individual fibres (75 Ni centres per nanometre; Supplementary Table 2) would imply that ∼45 NiBiD strands are present in parallel at each point within a fibre. Distributing these among the 10-20 nanoribbons observed per fibre (Fig. 1j), suggests that 2-5 NiBiD strands must be stacked in parallel to form a single nanoribbon (Fig. 4). A stacking of 2-5 layers at ∼0.35 nm distance, as also seen for stacked dimers of synthetic NBDT^33,34^, provides a nanoribbon height of 0.5-2 nm, which agrees well with the ribbon dimension as seen in STEM-HAADF (Supplementary Fig. 5d). The strand length (n value of Ni centers) of the Ni_n_(ett)_n+1_ complexes is not well constrained by the current data. Still, a monomeric form is not sufficiently conjugated to display resonance Raman scattering in the NIR region (see discussion above), and hence n>1. Likewise, the molar S/Ni ratio of 3.0-4.6 and spin concentration of 0.12 are only compatible with longer strand lengths n=20-40 (Extended Data Fig. 8). This suggests a stacking of long, planar, and extensively conjugated complexes, which would provide a low reorganization energy and high electronic coupling during electron transfer. The latter properties have previously been inferred to explain the high conductivities (> 100 S cm^-1^) of the fibre network^35^.

The alignment and stacking of NiBiD units into nanoribbons is further supported by polarized light microscopy of fibre skeletons, which show strong birefringence, thus suggesting that constituent components are preferentially oriented in parallel to the electron current (Fig. 3a-d). This polarized behaviour is consistent with previous Raman results, which show that the characteristic NiBiD modes display strong anisotropic scattering^16^. Angle-resolved Raman spectroscopy (−10 to 220°) confirms that the Ni-S stretching mode (367 cm^-1^) and the ring deformation mode (494 cm^-1^) only produce intense Raman signals upon excitation with light polarized in parallel (0°, 180°) to the conductive fibres (Fig. 3e). Polarized Raman spectroscopy additionally reveals that all NiBiD modes are highly polarized (depolarization ratio ρ < 0.25; Supplementary Table 7), thus suggesting that the 367 and 496 cm^-1^ modes constitute totally symmetric modes identical to those observed in NBDT complexes^36^. The NiBiD signals disappear almost completely when moving towards perpendicular polarization (90°) (Fig. 3f, Extended Data Fig. 9). Similar polarization effects are observed when exciting planar molecules aligned along their orientation axis with polarized light^37^. As the conductive fibres run in parallel to the longitudinal axis of the cable bacterium filaments, this indicates that the planar NiBiD units must be meticulously aligned with their long axis in parallel with the direction of the fibres.

**Fig. 3:**
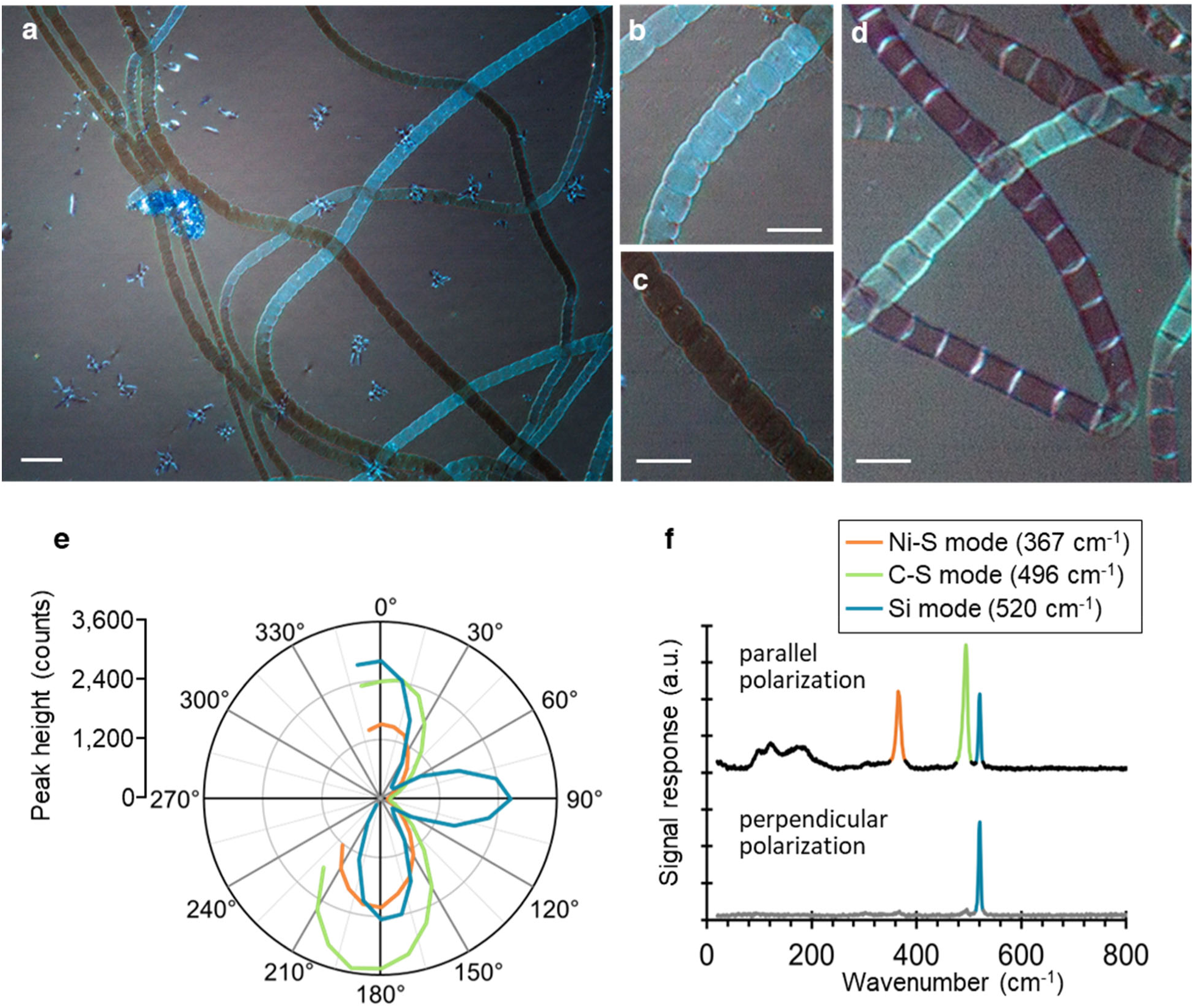
Polarization microscopy of cable bacteria. **a**, Polarized-light microscopy of native filaments of cable bacteria shows clear birefringence (40x objective). Scale bar: 10 µm. **b-c**, Enlarged sections of panel a displaying *Ca.* E. gigas filaments a. Filaments oriented at 45° (light) and 135° angles (dark). Scale bar: 5 µm. **d**, Polarized-light microscopy of fibre skeletons: the cell-cell junctions show the opposite behaviour as the fibres in the cell envelope. Scale bar: 5 µm. **e**, Fibre skeletons investigated by angle-resolved Raman spectroscopy show strong anisotropic scattering. Polar plot of the signal intensity (peak height) of the low-frequency modes (367 cm^-1^, 496 cm^-1^) of the NiBiD complex, using the Si mode from the substrate (520 cm^-1^) as a reference. **f**, Upon excitation with a polarization parallel to the fibre direction (0°, 180°), the NiBiD complex modes produce intense Raman signals. These signals disappear when the excitation polarization is perpendicular (90°).

### Discussion: A structural model for long-range conduction in cable bacteria

Our results establish a molecular basis for the highly efficient and long-range electron transport in the periplasmic fibres of cable bacteria. The mechanism is Ni-based, which aligns with a recent genomic intercomparison revealing that cable bacteria have a strongly expanded repertoire of nickel import and export enzymes, compared to their closest phylogenetic relatives^38^. Moreover, cable bacteria live in aquatic sediments, where porewaters feature elevated nickel concentrations (> 100 nM)^39^, thus ensuring nickel availability. Combining Raman, EPR, and XAS data with DFT calculations, we find that the Ni/S complex has a nickel(bisdithiolene) configuration, which has not been observed before in other enzyme cofactors. Molybdenum- and tungsten-containing proteins feature the molybdopterin cofactor, which has two adjacent thiolates coordinating the metal center^40,41^, while lactate racemase (LarA) contains the Ni-pincer cofactor, in which Ni is coordinated by two sulphurs, one carbon, and one imidazole (histidine)^29^. However, none of these cofactors feature the square planar -S_2_NiS_2_-geometry of NiBiD, nor do they display its extensive conjugation.

Our data suggest that the NiBiD complexes most likely have a Ni_n_(ett)_n+1_ oligomer configuration, and that these planar molecules are meticulously assembled into organo-metal “nanoribbons”, which form a set of parallel conduction channels in each fibre (Fig. 4). This intertwined ribbon structure bears a striking analogy to the braided copper wires used in the power cords of household appliances, and the purpose and advantages are likely similar. Cable bacteria are motile and make sharp turns^42^, all while the fiber network must remain fully functional and conductive. An entangled wire allows the conductor to be extremely flexible and malleable, but strong at the same time. An organo-metal framework configuration has not been previously seen in biology, but explains the unconventional electric properties that have been documented for cable bacteria fibres^6–9^. Nickel is a d8 electron transition metal and its dsp2 hybridization in NiBiD results in a square-planar configuration. The ett-ligands consist of π-conjugated fragments without rotatable single bonds, making the NBDT structure highly rigid, thereby facilitating the extension of electron conjugation along the oligomers and thus promoting charge transport^18^. Temperature-dependent characterization reveals that conductance of the fibre network remains high at temperatures <20 K, signifying quantum effects due to vibronic coupling^9^. In-plane vibrational modes of NiBiD are likely candidates to supply this electron-phonon coupling (Fig. 2a; Extended Data Fig. 4a).

**Fig. 4.**
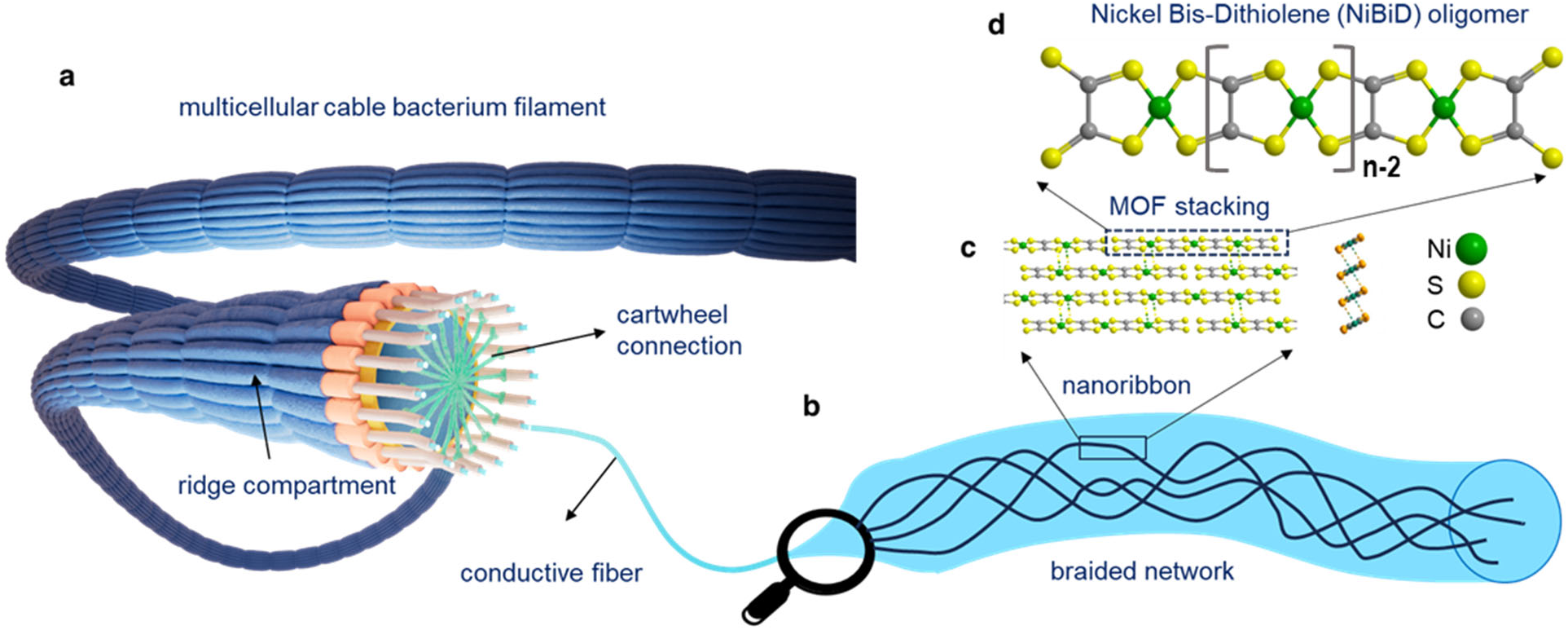
Structural model of the electron transport structure through the periplasmic fibre network in cable bacteria. **a**, A cable bacterium filament consists of a linear, unbranched sequence of cells. The surface is covered by parallel ridges that remain continuous along the filament. Each ridge compartment embeds a conductive fibre. Fibres are connected by a conductive cartwheel structure in the cell-cell interfaces. **b**, Each fibre embeds a entangled bundle of wavy conductive channels (nanoribbons). **c**, Each nanoribbon consists of planar NiBiD units that are axially aligned and stacked into a multilayer, one-dimensional organic framework. Side and cross-sectional views of a possible brick wall stacking are shown. **d**, Each planar NiBiD complex is an oligomer of nickel(ethene tetrathiolate) Ni_n_(ett)_n+1_, where n denotes the strand length (i.e. the number of linked Ni centres).

More generally, our findings uncover a mechanism through which electrical currents can be efficiently guided through an organic structure with a low metal content. The regular assembly of NiBiD oligomers into a long, one-dimensional transport channel appears to be key for fast electron relay. This configuration bears a striking similarity with synthetic 1D coordination polymers, like pNi(ett) and pNi(TTFtt)^19–21^, which can attain a high conductivity for a carbon-based material due to their extensive conjugation, electron delocalization, and overlapping π-systems^43^. Detailed electrochemical characterization has shown that fibre conductance exhibits similar organic-metal features, including a lack of redox signature, low reorganization energy, and high charge carrier density^8–9^.

The previously reported experimental values for the conductivity of the cable bacteria fibres (5 – 500 S cm^-1^) are calculated using the whole fiber cross-section (diameter 25 nm). If we restrict conductance solely to the nanoribbon network, then the estimated conductivity of a single nanoribbon (diameter 1.35 nm; 10-20 nanoribbons per bundle) amounts to 1×10^2^ – 2×10^4^ S cm^-1^. These are extremely high conductivity values for an organic structure, which exceed the reported values for synthetic polymeric Ni bis-dithiolenes (1-100 S cm^-1^) by two orders of magnitude. Accordingly, cable bacteria appear to have evolved a highly efficient form of organic conductivity, which could bring forward a novel design principle for sustainable electronic materials.

## Acknowledgements

The authors thank Peter-Leon Hagedoorn (Delft University of Technology) for guidance and discussion on EPR techniques, Alison Rodger (MacQuary University) for discussion on linear dichroism, Robin De Meyer (UAntwerpen) for assistance with SEM imaging, Saïd De Wolf (UAntwerpen) for helping out with nXRF image analysis, Benoit Desguin (UC Louvain) for providing LarA reference material, Nils Risgaard-Petersen for suggestions on the manuscript structure, and Jochen Blumberger (UCL) and Oliver Russell (UCL) for discussion on DFT modelling. We thank the staff at beamline ID16A-NI of the European Synchrotron Radiation Facility, beamline I20 at Diamond Light Source and the Penn State Materials Characterization Lab for experimental support. No editing services or AI-assisted technologies were used in preparation of the manuscript.

## Author contributions

Conceptualization: FJRM, HTSB

Culturing: SHM, JSG

Organic synthesis: YL, BCS

Raman spectroscopy: BS, HTSB, MW, GP, GN, KDW, FJRM

TEM microscopy: JAC, IP, JPT, TB, HR, FJRM

AFM microscopy: SHM

HAADF-STEM-EDX: NC, SHM, JAC, FJRM, SB, CT, JV

nXRF spectroscopy: BS, SHM, DK, SB

EPR spectroscopy: AC, MMR, BS, SHM, FJRM,

XAS spectroscopy: JSG, BS, SHM, GP, FJRM, MA, SH

DFT modelling: MA, MAZ

Data analysis: FJRM, BS

Structural model of cable bacteria: FJRM

Funding acquisition: FJRM

Project administration: FJRM

Supervision: FJRM

Writing – original draft: FJRM

Writing – review & editing: FJRM, BS, HTSB with input provided by all co-authors

## Additional information

Correspondence and requests for materials should be addressed to FJRM.

## Data availability

All source data related to this paper are available at a public repository (ZENODO) with reference (https://doi.org/10.5281/zenodo.10796906).

## Extended Data

**Extended Data Figure 1.**
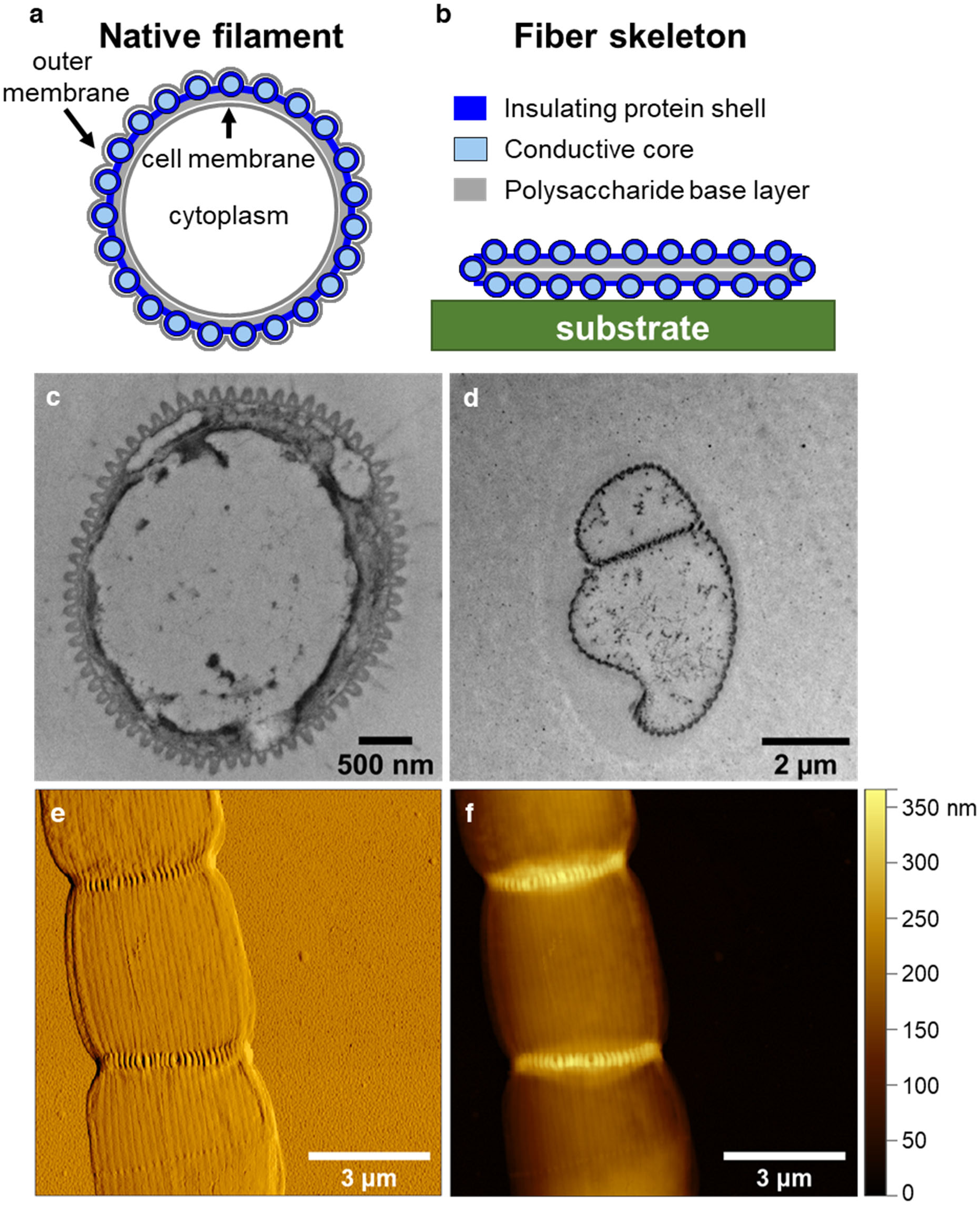
Fibre skeleton extraction procedure. **a-b**, Schematic of a cross-section across a native filament (a) versus a fibre skeleton (b). **c-d**, TEM images showing a cross-section of a resin-embedded native filament (c) and a fibre skeleton (d). **e-f**, AFM images of a fibre skeleton displaying the phase amplitude (e) and topography image (f).

**Extended Data Figure 2.**
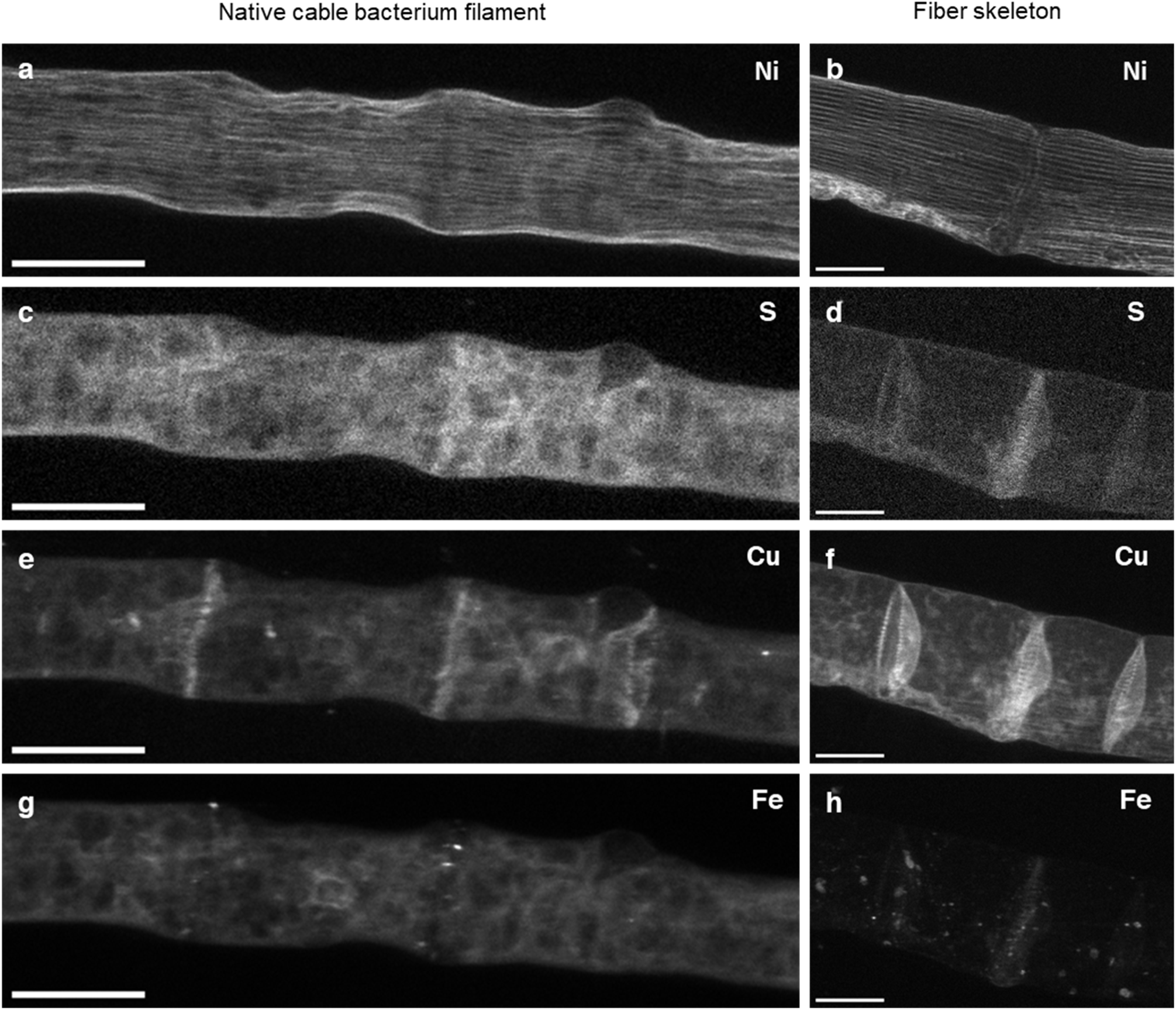
Synchrotron nXRF elemental maps of native cable bacterium filaments (a,c,e,g) and fibre skeletons (b,d,f,h). Gray values reflect area densities (ng mm^-2^). Elements are indicated in top right corner of each panel. Panels b and f are a grayscale replication of Fig. 1e and Fig. 1f in the main text, respectively. Scale bars: 2 µm.

**Extended Data Figure 3.**
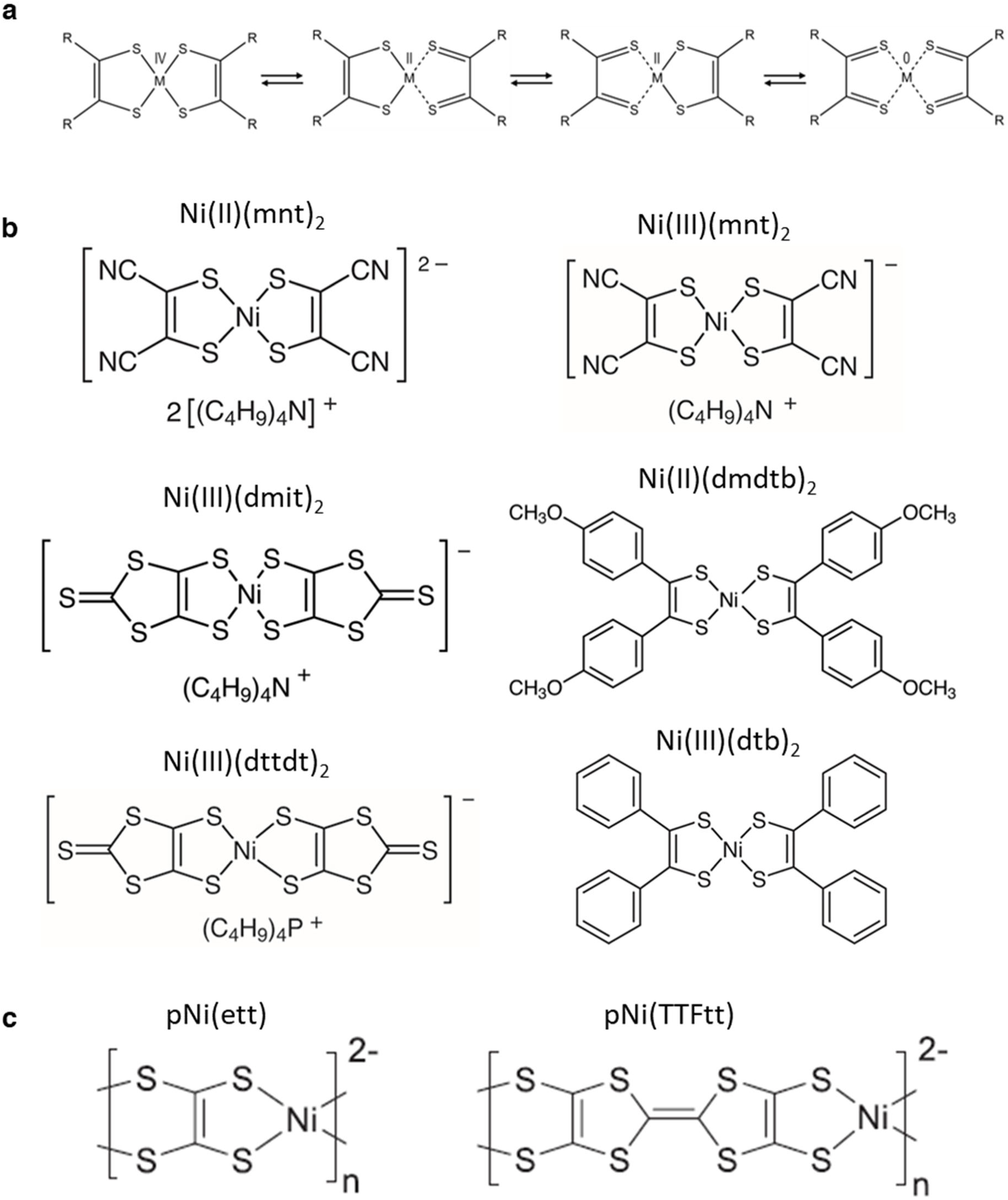
Structural formula of the synthetic NBDT compounds used as reference compounds in spectroscopic analysis. **a**, Generic structure of a metal bis-dithiolene. R represents a generic substituent on the dithiolene ligand. **b**, Structure of six monomeric NBDT reference compounds used as reference. **c**, Structure of the polymeric NBDT reference compounds pNi(ett) and pNi(TTFtt). See Extended Data Table 1 for nomenclature.

**Extended Data Figure 4.**
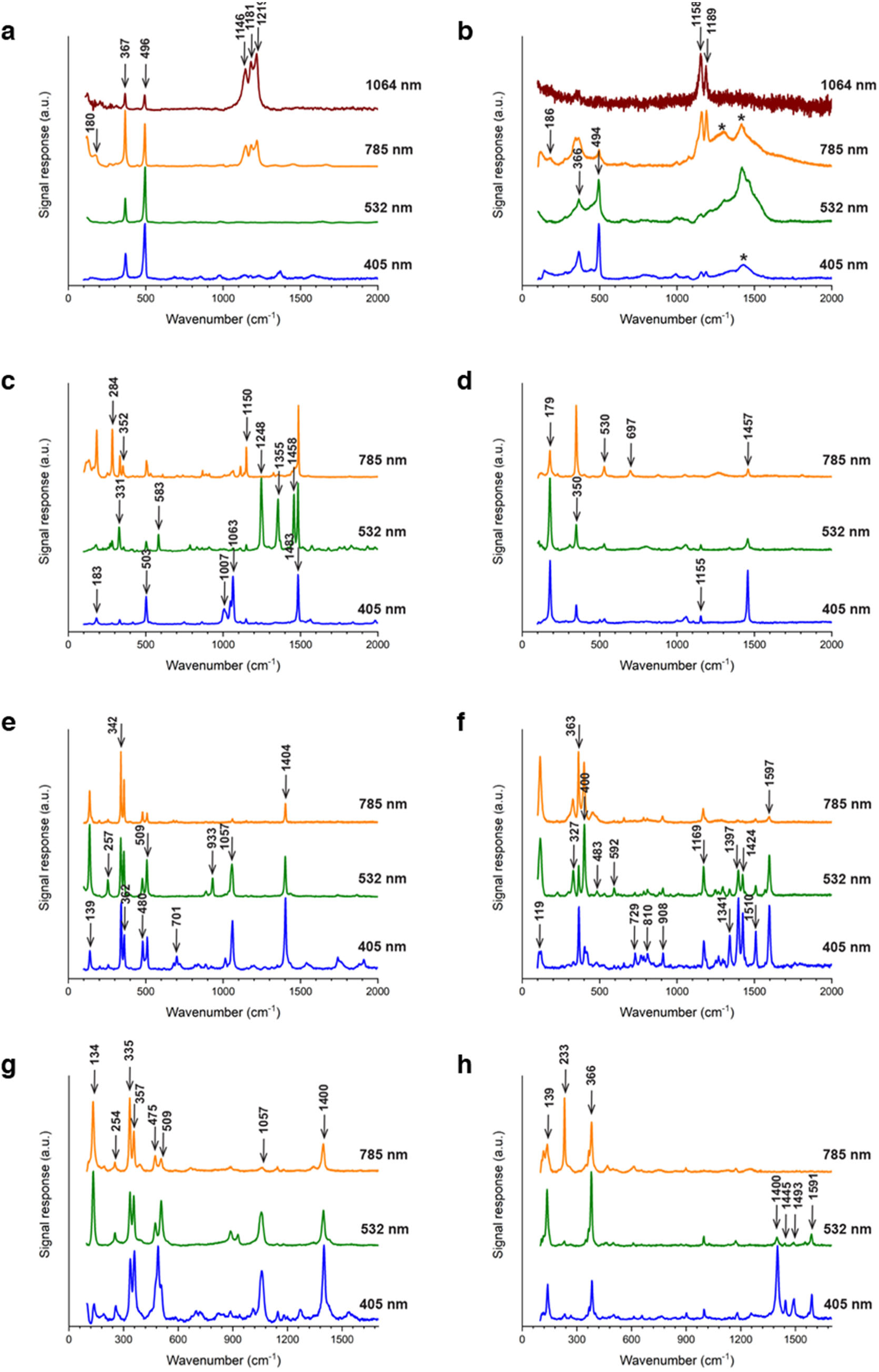
Raman spectra of cable bacterium fibre skeletons and NBDT reference compounds. **a**, Fibre skeletons of cable bacteria. **b**, pNi(ett) **c**, Ni(II)(mnt)_2_ **d**, Ni(III)(mnt)_2_ e, Ni(III)(dmit)_2_ **f**, Ni(II)(dmdtb)_2_ **g**, Ni(III)(dttdt)_2_ **h**, Ni(II)(dtb)_2_. See Extended Data Table 1 for nomenclature and Extended Data Fig. 3 for molecular structures.

**Extended Data Figure 5.**
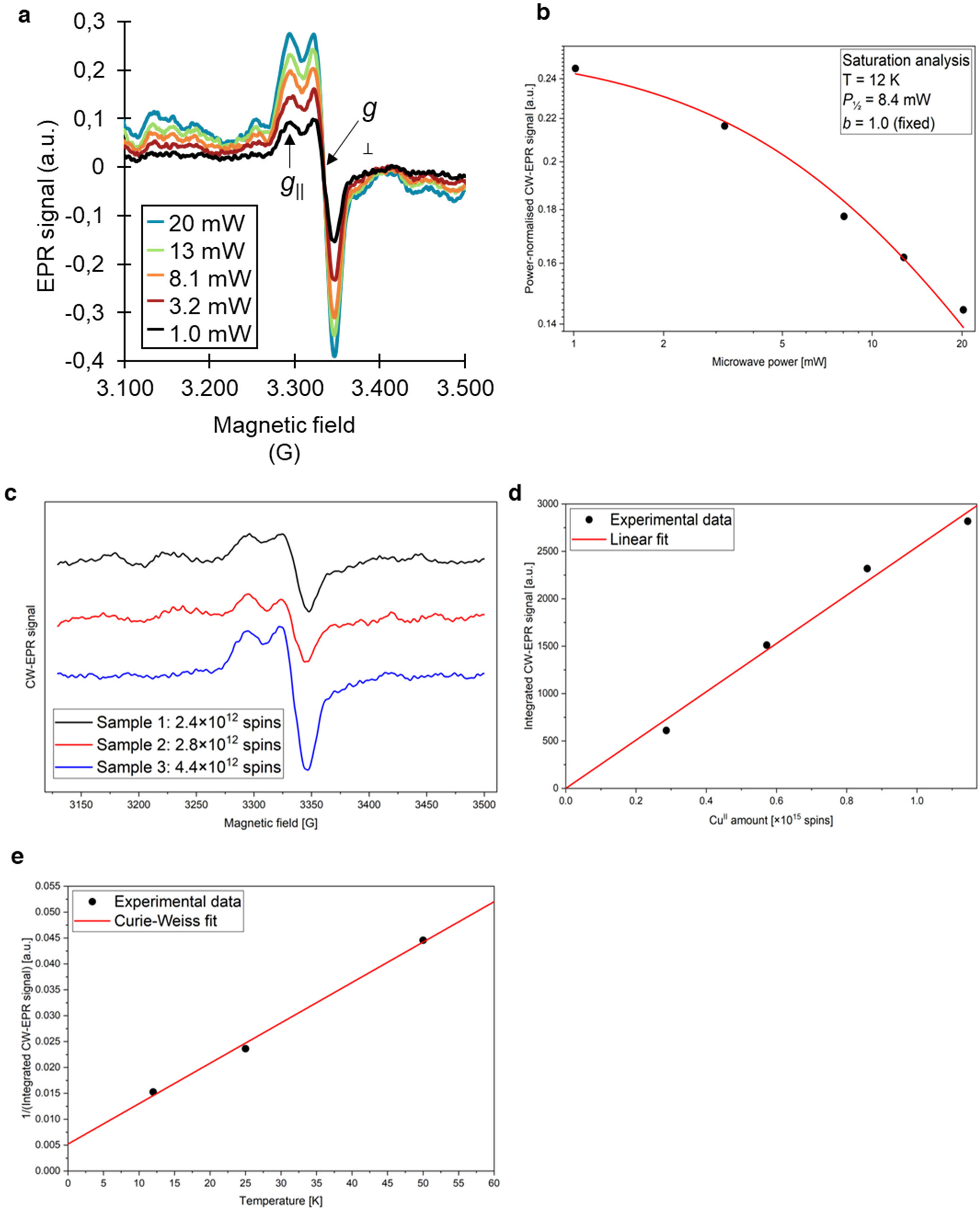
Electron Paramagnetic Resonance (EPR) spectroscopy of native cable bacterium filaments. **a**, Microwave power dependence of the observed EPR signal at 12 K. **b**, Power saturation plot (logarithmic axes) based on the data shown in panel a. The *Power-normalised CW-EPR signal* (y axis) is calculated as the peak-to-peak amplitude of the *g*_⊥_ component divided by the square root of the corresponding microwave power. The EPR signal is only marginally saturated even at high microwave power (20 mW) and low temperature (12 K), consistent with a relatively fast-relaxing species. The equation used for the fitting curve and the meaning of the parameters *P*_1/2_ and *b* are described in the Methods section. An accurate determination of these parameters is not possible as full saturation could not be achieved in the explored microwave power range. The inhomogeneity parameter *b* (1 ≤ *b* ≤ 3) gave the best agreement with the experimental data for the value b = 1. **c**, The EPR signals obtained from 3 biological replicates are similar. All displayed traces were measured with a microwave power of 1.0 mW. **d**, The number of spin centres in each sample was estimated via a Cu^II^ calibration curve as described in the Methods section. **e**, The reciprocal of the double integral (S) of the CW-EPR signal, recorded at 0.2 mW microwave power and 20 G field modulation amplitude, is plotted against the temperature T to investigate the magnetic behaviour of the samples. The solid line is the fit (1/S vs. T) of the experimental data points to the Curie-Weiss relation, S = C/(T-θ).

**Extended Data Figure 6.**
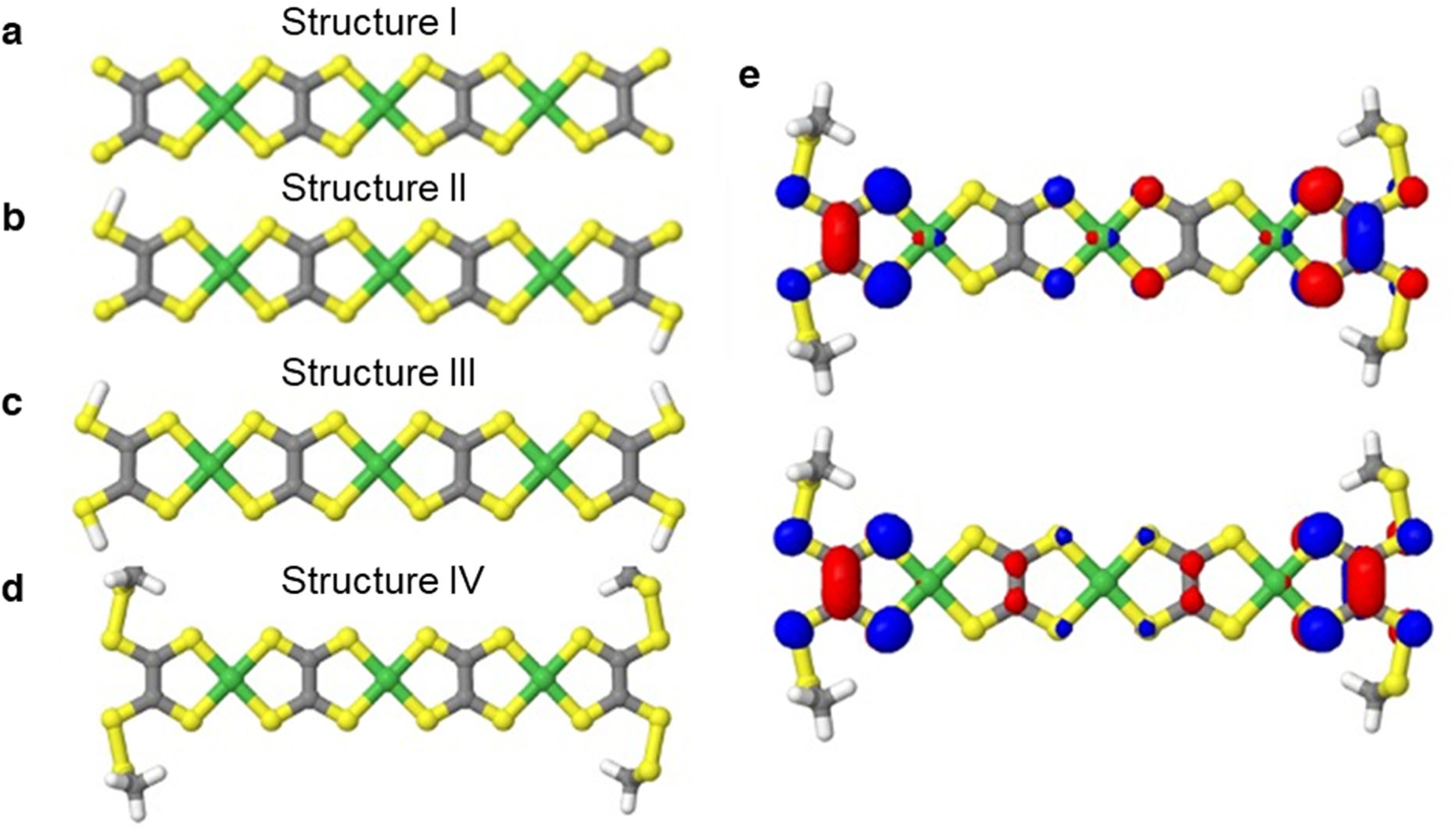
DFT optimized geometries of oligomeric Ni_3_(ett)_4_ oligomers as representative models for NiBiD. All four structures contain three Ni atoms and four C_2_S_4_ tetrathiolate ligands but have distinct end-capping groups. **a**, Di-anionic oligomer with only terminal sulphurs. **b**, Charge-neutral oligomer where two of the terminal sulphurs are capped with hydrogen atoms. **c**, Charge-neutral oligomer where all four terminal sulphurs are capped with hydrogen atoms. **d**, Charge-neutral oligomer where the terminal sulphur atoms have formed S-S bonds (crosslinking with thiol groups of cysteine residues is mimicked by addition of -SCH_3_ groups). **e**, Plots displaying the two highest singly occupied orbitals of Structure IV optimized as an open shell singlet showing the unpaired electrons localizing on the sulphur and carbons atoms of the terminal ligands with minimal contribution of the nickel atoms.

**Extended Data Figure 7.**
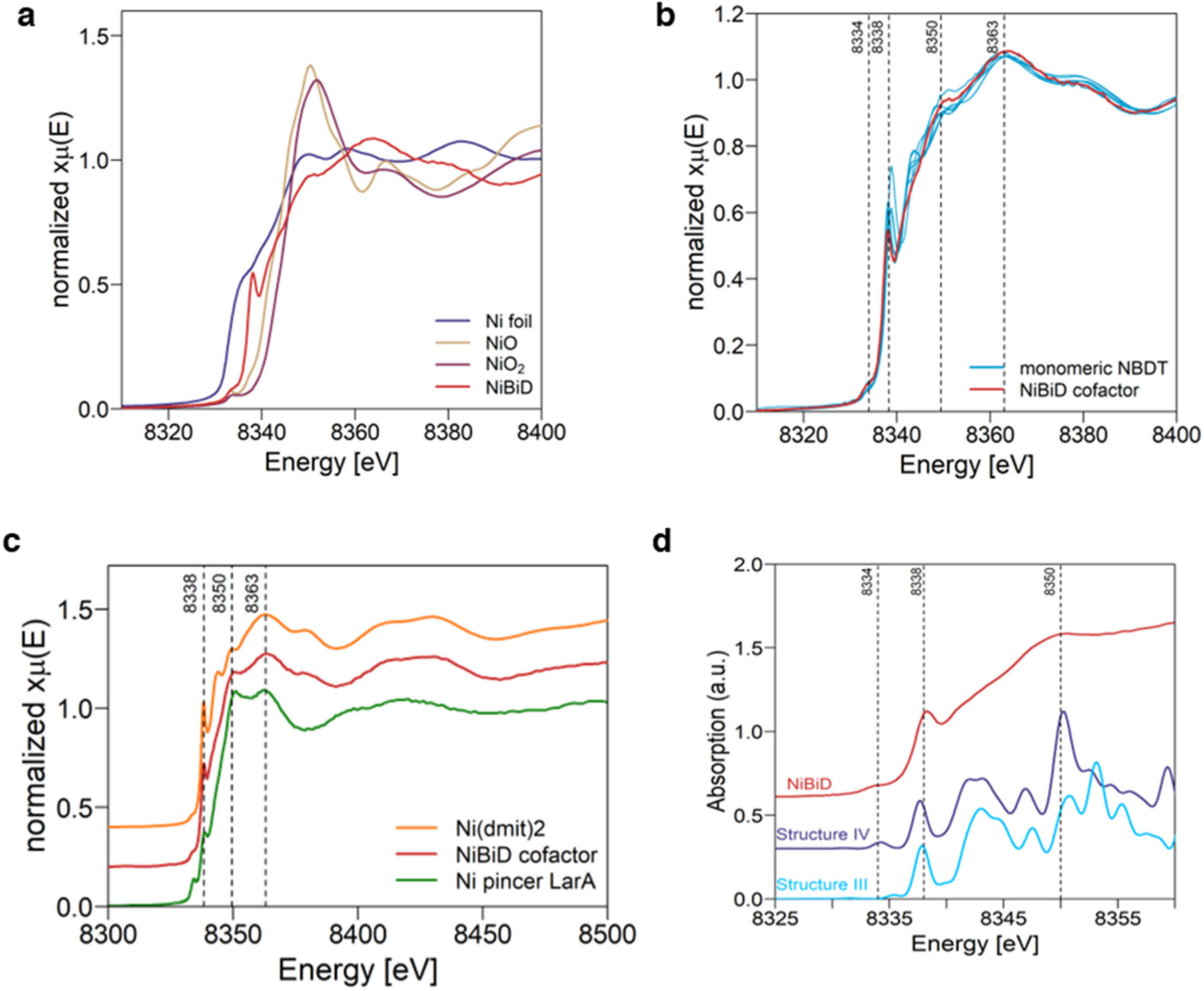
X-ray absorption spectra at the Ni K-edge. **a**, The Ni K-edge of the NiBiD complex in native filaments has a different shape and position than the inorganic references. **b**, The Ni K-edge XANES spectrum of the native filaments containing the NiBiD complex is compared to that of 6 monomeric nickel bis(1,2-dithiolene) complexes (list in Extended Data Table 1). The spectra are highly similar, indicating NiBiD has a clear NBDT character. **c**, The Ni K-edge XAS spectrum of native cable bacterium filaments containing the NiBiD complex is compared to that of the monomeric NBDT compound Ni(III)(dmit)_2_ and the LarA enzyme containing the Ni pincer. Spectra in panels c and d are offset by 0.2 for clarity. **d**, TD-DFT simulations of Ni_n_(ett)_n+1_ structures III and VI in Extended Data Fig. 6 compared to the experimental NiBiD spectrum. Structure IV shows a similar pre-edge as NiBiD. Spectra are off-set for clarity. a.u. = arbitrary units.

**Extended Data Figure 8.**
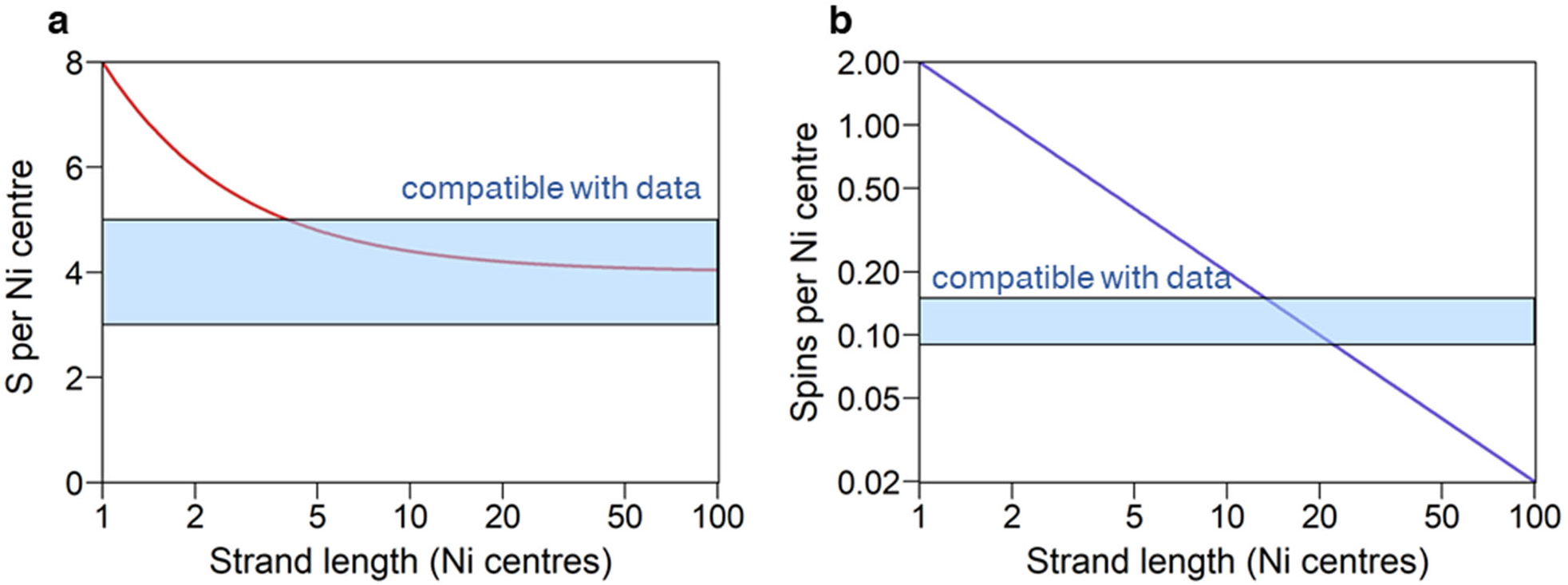
Experimental constraints on the NiBiD strand length. **a**, The red solid curve specifies the theoretical S/Ni ratio in Ni_n_(ett)_n_, i.e., (4n+4)/n where n is the strand length (number of Ni centres). The molar S/Ni ratio range of 3-4.6 for NiBiD as obtained from nXRF and STEM-HAADF-EDX provides a constraint on the strand length (blue rectangle). **a**, The blue solid curve specifies the theoretical spin concentration in Ni_n_(ett)_n_, assuming its an open shell triplet with two spins per molecule, i.e., 2/n where n is the strand length. The experimental range of 0.12±0.03 as obtained by EPR is indicated by the blue rectangle.

**Extended Data Figure 9.**
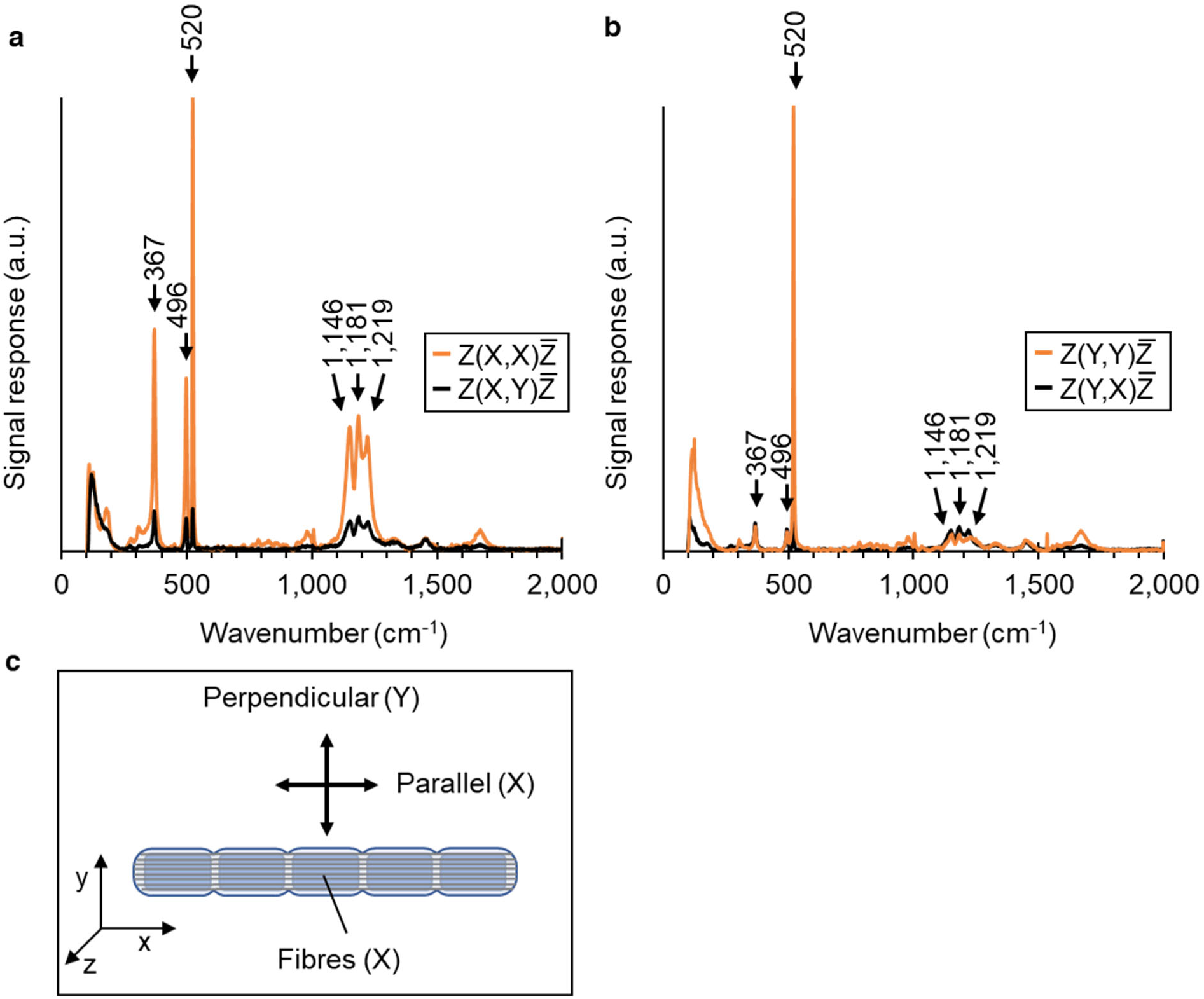
Parallel and cross-polarized Raman spectra of fibre skeletons. **a**, Spectra recorded with the laser polarization in parallel (X) to direction of the conductive fibres (X). **b**, Spectra recorded with the laser polarization perpendicular (Y) to the direction of the conductive fibres (X). All Raman spectra were recorded with the 785 nm excitation laser. The peak at 520 cm^-1^ arises from the Si substrate. **c**, Schematic of the Polarized Raman spectroscopy sample setup. Cable bacteria were positioned on the substrate so that conductive fibres were oriented in the x-direction. Laser light, incoming via the out-of-plane z-axis, was polarized in the X-(parallel) or Y-direction (perpendicular) to record Raman spectra with an analyser.

**Extended Data Table 1.** List of nickel bis(dithiolene) reference compounds used in Raman and XAS spectroscopy. The associated molecular structures are displayed in Extended Data Fig.3. Type refers to monomeric (“M”; 1 Ni centre) or polymeric (“P”; >1 Ni centre). Monomeric compounds were all commercially obtained from TCI Chemicals with catalogue number provided (Cat. Nr.). Polymeric compounds were synthesized following the procedure in Ref 20.

| Type | Code | Cat. Nr. | R Ligand | Name |
| --- | --- | --- | --- | --- |
| M | Ni(II)(mnt)<sub>2</sub> | B1371 | -C≡N | bis(tetrabutylammonium)<br>bis(maleonitriledithiolato) nickel(II) |
| M | Ni(III)(mnt)<sub>2</sub> | T1415 | -C≡N | tetrabutylammonium<br>bis(maleonitriledithiolato) nickel(III) |
| M | Ni(III)(dmit)<sub>2</sub> | T1272 | S<sub>2</sub>C=S | tetrabutylammonium bis(1,3-dithiole-2-thione-4,5-dithiolato) nickel(III) |
| M | Ni(II)(dmdtb)<sub>2</sub> | B4360 | -Phe-O-CH<sub>3</sub> | bis[4,4'-dimethoxy(dithiobenzil)] nickel(II) |
| M | Ni(III)(dttdt)<sub>2</sub> | T1416 | S<sub>2</sub>C=S | tetrabutylphosphonium bis(1,3-dithiole-2-thione-4,5-dithiolato) nickel(III) |
| M | Ni(II)(dtb)<sub>2</sub> | B1350 | -Phe | Bis(dithiobenzil) nickel(II) |
| P | pNi(ett)<sub>2</sub> | synthesized | -SH | poly-nickel (ethene tetrathiolate) |
| P | pNi(TTFtt)<sub>2</sub> | synthesized | S<sub>2</sub>C=CS<sub>2</sub> | poly-nickel (TetraThiaFulvalene<br>tetrathiolate) |

**Extended Data Table 2.** Spin quantification. Three replicate samples composed of native cable bacterium filaments were analysed. The column “#EPR spins” shows the spin number as derived from the calibration curve (Extended Data Fig. 5d). The Filament Length (L_fil_) represents the total length of cable bacterium filaments in each EPR tube as determined by the imaging procedure (see Methods). The “Fibre length” is calculated as the L_fibre_ = L_fil_ x N_fibre_, where N_fibre_ = 68 is the number of fibres as determined by TEM and AFM. The number of Ni centres in the sample is determined as #Ni = L_fibre_* X_Ni_ with X_Ni_ = 75 denoting the number of Ni centres per nm of fibre (as determined from nXRF imaging – see Supplementary Table 3). The spins per Ni centre is finally calculated as the ratio (#EPR spins)/(#Ni).

| Sample | #EPR spins | Filament<br>Length (mm) | Fibre<br>Length (nm) | #Ni | Spins per Ni |
| --- | --- | --- | --- | --- | --- |
| 1 | $2.4 \times 10^{12}$ | 4758 | $3.24 \times 10^{11}$ | $2.41 \times 10^{13}$ | **0.10** |
| 2 | $2.8 \times 10^{12}$ | 3660 | $2.49 \times 10^{11}$ | $1.86 \times 10^{13}$ | **0.15** |
| 3 | $4.4 \times 10^{12}$ | 8260 | $5.62 \times 10^{11}$ | $4.19 \times 10^{13}$ | **0.10** |

## EXPERIMENTAL MODEL DETAILS

### Cable bacteria cultivation

Cable bacterium filaments were produced by enrichment culturing and clonal culturing. The natural sediment used for culturing was collected from an intertidal creek within Rattekaai salt marsh (51.4391°N, 4.1697°E; The Netherlands). Enrichment culturing of cable bacteria was performed as described previously^44^. Briefly, natural sediment was first sieved to remove debris, plant material, and fauna (1.4 mm stainless steel mesh). Homogenized sediment was subsequently packed into transparent PVC core liners (inner diameter 36 mm, height 100 mm) with a rubber stopper at the bottom. Sediment cores were submerged in plastic containers containing artificial seawater (salinity 30, Instant Ocean) that was continuously aerated, and incubated at a constant temperature (20 °C) in the dark. After 4-6 weeks, a dense population of thick (∼ 4 µm diameter) cable bacteria developed in the sediment cores of enrichment culture.

Because one starts from a natural sediment, enrichment cultures can contain multiple strains of cable bacteria. Therefore, we created a so-called clonal culture that contained only a single strain of cable bacteria. This clonal culture (strain JX3-16) was attained from the same Rattekaai sediment source via transfer of individual filaments to autoclaved sediment cores as described in detail in Ref.^38^. To confirm clonality (i.e. the presence of only a single cable bacterium strain), we performed V3-V4 16S rRNA amplicon sequencing^45^. The clonal culture was maintained in the lab by regularly transferring a small amount of the clonal enrichment culture as inoculum to freshly autoclaved sediment cores. The JX3-16 strain belongs to the species *Candidatus* Electrothrix gigas^45^.

Individual cable bacteria filaments were harvested from the sediment cores derived from the cultures. Hereto, a small portion of sediment was deposited on a glass microscopy cover slip. Custom-made glass hooks were used to isolate cable bacterium filaments from the sediment and handle them under a stereomicroscope. Filaments were washed six times by transferring them between droplets (∼ 20 µl) of milliQ water (mQ), thus removing attached particles and salts, until clean filaments were obtained (further referred to as “native filaments”).

During the initial phase of the study, cable bacterium filaments were retrieved from enrichment cultures with *Ca.* E. gigas and used for EPR and nXRF analysis. Subsequently, when the JX3-16 clonal culture became available, filaments were solely retrieved from this clonal culture for analysis (Raman, XAS, HAADF-STEM-EDX). To verify consistency, we compared the morphology of the filaments retrieved from the enrichment and clonal cultures via TEM cross-sections at various time points. The thick filaments initially retrieved from enrichment cultures displayed a similar size and morphology, and also had the similar number of ridge compartments as the JX3-16 strain, thus indicating that alle filaments investigated belonged to the same species i.e., *Candidatus* Electrothrix gigas^44^.

### Fibre skeleton extraction

Through sequential extraction, the conductive fibre network was isolated from the cell envelope of individual filaments, as previously described^5,6,15^. This extraction procedure removes the membranes and cytoplasm, thus producing a so-called “fibre skeleton” (Extended Data Fig. 1). Hereto, native filaments were deposited for 10 minutes in a droplet (∼ 20 µl) of 1% (w/w) sodium dodecyl sulphate (SDS) to break down the cell membranes and release the cell content. After 10 minutes, filaments were sequentially rinsed 6 times in fresh mQ droplets to remove cell remains and wash off excess SDS. Next, filaments were submerged for 10 minutes in a droplet (∼ 20 µl) of 1 mM ethylenediaminetetraacetic acid (EDTA), adjusted to pH 8, to remove residual SDS. Filaments finally underwent a final 6 – 10 washes in fresh mQ droplets to obtain fibre skeletons for subsequent analysis. The quality of fibre skeleton extraction was monitored with Raman microscopy, TEM and AFM imaging (see Extended Data Fig. 1 and Supplementary Text).

Fibre skeletons were further fragmented into subcomponents through sonication, thus producing individual fibres and pieces of the cartwheel (further referred to as “fractionated fibre skeletons”). To this end, fibre skeletons were transferred to 200 µL of mQ in an Eppendorf 1.5 mL vial. The sonicator (Q125 probe sonicator, Qsonica) operated at 20% amplitude for 10 minutes in a cyclic fashion (30 seconds on alternated by 30 seconds off). The vial was cooled with ice while sonicating.

## METHOD DETAILS

### Atomic Force Microscopy

Fibre skeletons were transferred onto a 50 nm gold coated silicon wafer (Platypus Technologies) in a drop of mQ water and subsequently air-dried. The wafer was securely fixed with double sided carbon tape to a 12 mm diameter stainless steel metal disc. AFM images were acquired in tapping mode with a XE-100 scanning probe microscope (Park Systems) equipped with an aluminium SPM probe with a tip radius < 10 nm (AppNano ACTA-200), resonant frequency of 200-400 kHz, and a nominal spring constant of 13–77 N/m. Topographic and amplitude data were recorded and processed with the Gwyddion software^46^.

### Polarized-light microscopy

Polarized-light microscopy was performed on native cable bacterium filaments retrieved from a natural sediment incubation of Rattekaai sediment, containing *Ca.* E. gigas but also from thinner species of cable bacteria (Fig. 3a-d). Images were collected with an Olympus BX50 microscope with the polarizer (at 90°) and analyser (at 0°) crossed. Images were captured with a Canon 6D full-frame digital camera. Adobe Camera Raw was used to set image contrast and brightness with curves and to edit colour balance.

### Scanning Electron Microscopy

SEM images were acquired using a FEI Quanta 250 FEG environmental scanning electron microscope (Thermo Fisher Scientific). Before imaging, filaments were coated with an Au layer via sputtering (Leica EM ACE600). Images were collected in Secondary Electron (SE) mode using an accelerating voltage of 20 kV and a working distance of 10 mm.

### Transmission Electron Microscopy

Fibre skeletons and fractioned fibre skeleton samples were pipetted onto a 200 mesh Formvar-coated copper grid (Micro to Nano) and allowed to air-dry on the grids. The samples were stained with 2% uranyl acetate for 1 minute. After air-drying, the grids were examined with two TEM instruments: a FEI Tecnai G2 Spirit BioTWIN operating at 120 kV, or a JEOL 1400 TEM equipped with LaB6 filament and TVIPS F416 CCD camera also operating at 120 kV.

To arrive at resin embedded cross-sections, native cable bacterium filaments and fibre skeletons were incubated for 15 minutes in freshly prepared fixative (4% paraformaldehyde, Sigma-Aldrich) in 1× phosphate-buffer saline (PBS; 137 mM NaCl, 2.7 mM KCl, 10 mM Na_2_HPO_4_, 1.8 mM KH_2_PO_4_, pH 7.4) at room temperature. Fixative removal involved two 3-min washes in 1× PBS, followed by two 3-minute washes in 0.1× PBS and a final rinse in mQ water. Samples were then embedded in a thin layer of 2% low melting point agarose (Fisher Bioreagents). After solidifying, agarose-embedded specimens were reshaped into small rectangles and subjected to secondary fixation in 2% paraformaldehyde in 0.1 M phosphate buffer (pH 7.4), washed with 0.01 M phosphate buffer (pH 7.4), and dehydrated using a graded ethanol series (50%, 70%, 90%, 95%, and three changes of 100% ethanol for 20 minutes each). Infiltration of Unicryl resin in the agarose blocks was accomplished by incubation in 30% resin in ethanol for 1 hour, followed by at least three changes with fresh 100% resin. Subsequently, samples were submerged in fresh Unicryl resin and cured under UV light for 60 h at 4°C followed by 72 h at room temperature. The resulting Unicryl blocks were sectioned using an ultramicrotome equipped with a diamond knife, yielding 50 nm thick sections. Thin sections were then transferred onto Au TEM grids (Electron Microscopy Sciences). Next, sections were stained with 2% uranyl acetate and lead citrate for 1 minute, washed, and dried before being examined with transmission electron microscopy (TEM). Thin sections were imaged with a FEI Tecnai G2 Spirit BioTWIN operating at 120 kV.

### Synchrotron nano-X-ray Fluorescence elemental mapping

Native cable bacteria filaments, intact fibre skeletons, and fractioned fibre skeleton samples were subjected to nXRF imaging. Samples were deposited on silicon nitride membranes (Silson). A droplet of ultrapure water (Optima™ LC/MS Grade, Thermo Fisher Scientific) was deposited over the sample and blotted with paper to leave a thin layer of liquid. Next, the sample was plunge frozen in liquid ethane at 93 K (Leica EM GP), transferred to a bath of liquid nitrogen (Leica EM VCM), and inserted in a vacuum cryo-transfer shuttle system (Leica EM VCT500). During transfer, the shuttle was kept at a temperature between 120 and 113 K. The cryo-transfer system was then connected to the experimental chamber of the ID16A Nano-Imaging Beamline (European Synchrotron Radiation Facility, Grenoble, France) to introduce the sample on the sample stage, which is kept in cryogenic condition (100 K) to reduce the radiation damage on the sample during X-ray fluorescence mapping. A light microscope focused on the sample was used to locate the sample on the membrane and to define the scanning region. The X-ray beam was operated at a constant energy of 17 keV and focused to a spot size of 35 nm. The nXRF maps were obtained by scanning the beam over the fibre skeletons in 10-25 nm steps and recording the X-ray fluorescence signal emitted from each sample pixel with two silicon drift detectors (16 element Ardesia and 7 element Vortex ME-7) positioned perpendicular to the X-ray beam^47^. The elemental areal mass concentration was calculated using the Fundamental Parameters (FP) approach implemented in PyMca software package^48^ and a reference standard material containing elements of certified concentration (RF7-200-S2371, AXO Dresden) with uniform mass depositions in the range of ng/mm² (1-3 atomic layers).

### STEM-HAADF-EDX imaging

Scanning Transmission Electron Microscopy (STEM) imaging was performed in High-Angle Annular Dark-Field (HAADF) mode, combined with Energy Dispersive X-Ray (EDX) mapping, on a Tecnai Osiris (Thermo Fischer Scientific), equipped with a high brightness X-Feg electron, an analytical pole piece (6 mm pole piece gap), a 4 quadrant Super X detector and a Gatan US1000XP CCD camera. The microscope was operated at 200 kV with a beam current of ∼150-200 pA. Image analysis was performed in the Quantax software, with additional postprocessing in ImageJ and R.

### Nickel bis(dithiolene) reference compounds

Six monomeric (single Ni centre) nickel bis(1,2-dithiolene) complexes were obtained as powders from TCI Europe n.v. Ni(II)(mnt)_2_, Ni(III)(mnt)_2_, Ni(III)(dmit)_2_, Ni(II)(dmdtb)_2_ Ni(III)(dttdt)_2_ and Ni(II)(dtb)_2_ - catalogue numbers provided in Extended Data Table 1. Two polymeric nickel bis(1,2-dithiolene) complexes, Poly-nickel(ethene tetrathiolate) (pNi(ett)) and Poly-nickel(TetraThiaFulvalene tetrathiolate) (pNi(TTFtt)), were synthesized as powders according to the protocol described in Liu et al.^20^. The chemical formulas are listed in Extended Data Table 1 and the associated molecular structures are depicted in Extended Data Fig. 3. All NBDT compounds were stored under dark and anaerobic conditions (N_2_ glovebox) until further use. LarA was obtained by the procedure described in Desguin et al.^29^.

### Raman microscopy

For multi-wavelength Raman microscopy, reference compounds included six monomeric NBDT compounds (Extended Data Fig. 3b, Extended Data Table 1) and the two polymeric NBDT compounds pNi(ett) and p(TTFtt) (Extended Data Fig. 3c). Fibre skeletons and NBDT powders were deposited on Si wafers coated with 50 nm gold or small pieces of aluminium foil attached to a microscopy slide. Raman spectra were collected using a Renishaw inVia™ Qontor® microscope equipped with four Renishaw solid state lasers (405 nm, 532 nm, 785 nm, 1064 nm). We used a 20x (NA 0.50) microscopy objective and confocal aperture of 65 µm to focus incident laser light on the samples and capture Raman scattered photons. Two electrically cooled (−70 °C) detectors were used to detect Raman scattered photons in the visible range (Renishaw Centrus Charge Coupled Device) and near-infrared range (Andor Indium Gallium Arsenide photodiode). Spectrum acquisition parameters, including grating, acquisition time, and laser power were optimized to ensure optimal signal response and spectral resolution (± 1 cm^-1^), and are summarized in Supplementary Table 4. Raman spectra were processed using the LabSpec6 software (version 6.6.1.11; Horiba). Processing was limited to cosmic ray removal and baseline correction. Baselines were removed by fitting a polynomial to the acquired spectrum. Finally, processed spectra were averaged to improve the signal-to-noise ratio.

For angle-resolved Raman spectroscopy, fibre skeletons were deposited onto a Si wafer coated with 50 nm gold (Platypus Technologies). Spectra were acquired using a Horiba LabRam HR Evolution confocal Raman microscope equipped with a 633 nm HeNe laser (ThorLabs HRP-350-EC). An analyser was placed in front of the spectrometer such that parallel polarized Raman scattered light was recorded as a function of the incident angle. The optical components of the Raman system included a 100x objective (0.90 NA), a confocal hole aperture of 50 μm, and 600 gr/mm grating. A back-illuminated deep-depleted Si array (1,024 x 512 pixel) detector, calibrated using the Raman response of a single crystal Si sample, was used to detect Raman scattered light. Spectra were collected by focusing the laser spot, delivering approximately 12 mW of laser power during 1 second, on the middle of the fibre skeletons. An automated ½ waveplate in the common path controlled the polarization of the excitation laser, which was rotated from −10° to 220° with respect to the direction of the periplasmic protein fibres in the filaments.

Polarized Raman microscopy was done using the 785 nm excitation laser of the Renishaw inVia™ Qontor® microscope. A λ/2 waveplate was put in the optical path to change the incident polarization between horizontal (X) and vertical (Y) polarization. An analyser kit, consisting of a λ/2 waveplate and a polarizer, was used to examine the polarization of the Raman scattered light. This setup enabled us to test 4 polarization conformations: Z(X,X)Z̅, Z(X,Y)Z̅, Z(Y,X)Z̅, and Z(Y,Y)Z̅. Spectra were collected for fibre skeletons that were oriented horizontally (X) on the Si wafer (Extended Data Fig. 8c). The wafer was oriented with the crystal axes parallel to the polarizations of the incident laser light. This way, the 520 cm^-1^ Si-mode functioned as a well-characterized, polarized reference mode. The excitation laser delivered 5 mW of laser power during 30 seconds to the sample. Per conformation, 10 spectra were recorded at different spots in the fibre skeleton. After processing, spectra were averaged to obtain the final spectra shown in Extended Data Fig. 8a,b. The depolarization ratio (ρ) of the NiBiD complex modes was calculated (Supplementary Table 7) as the ratio of the intensity of perpendicular versus the parallel component of the Raman signal (Z(X,Y)Z̅ /Z(X,X)Z̅).

### X-band Electron Paramagnetic Resonance (EPR) spectroscopy

Native cable bacterium filaments were individually retrieved from the enrichment incubation, cleaned, and gathered in a dense clump (> 1000 filaments) in droplet of mQ water on a microscope slide. This clump was then transferred to a Suprasil EPR tube (100 mm length, 1.1 mm inner diameter (ID), 1.6 mm outer diameter (OD); Wilmad Labglass WG-222T-RB) filled with mQ water. The EPR tube was inserted in a 15 mL conical centrifuge tube together with filler material to avoid breaking the tube during centrifugation, and the tube assembly was centrifuged for 5 minutes at 2,500 g. As a result, the filament clump migrated to the bottom of the EPR tube and formed a small pellet. The supernatant was removed from the EPR tube using a syringe with a long needle until ±1 mm of water remained, and subsequently, the EPR tube was flash frozen in liquid nitrogen and stored at 77 K until further use. Three replicate EPR tubes with sample were prepared according to this procedure.

EPR measurements were performed on a Bruker EMX X-band continuous wave (CW) EPR spectrometer equipped with a Bruker ER 4122SHQE super-high-sensitivity resonator and a closed-circuit Helium cryostat (Cryogenic Ltd) controlled by a Model 350 temperature controller (Lake Shore Cryotronics, Inc.) and a Cernox sensor (Lake Shore Cryotronics, Inc.). The magnetic field was calibrated using 2,2-diphenyl-1-picrylhydrazyl (DPPH) as a standard (*g* = 2.0036)^50^. Immediately before measurement, the Suprasil EPR tube containing the sample was loaded inside a clear fused quartz (CFQ) tube (2 mm ID, 3 mm OD, Wilmad Labglass 705-SQ-250M). The latter was evacuated to a final pressure of approximately 1 mbar using a 3 mm NMR tip-off manifold (Wilmad Labglass) to prevent condensation of liquid oxygen inside the EPR tube during the low-temperature measurements. This whole procedure was performed under liquid Nitrogen to prevent the biological sample from thawing. The assembly was loaded in the pre-cooled cryostat so that the pellet containing the cable bacteria is positioned at the centre of the EPR resonator. For each of the three replicates CW-EPR spectra were recorded at 12 K using microwave powers of 20.2 mW, 12.8 mW, 8.0 mW 3.2 mW and 1.0 mW (unless indicated otherwise), a field modulation of 20 G at 100 kHz, a magnetic field sweep rate of 13.3 G/s, a resolution of 1.02 points/G, a conversion time of 73.44 ms and a time constant of 20.48 ms. The microwave frequency was ≈9.37 GHz. Background spectra from an empty CFQ quartz tube (2 mm ID, 3 mm OD) were recorded under the same experimental conditions. The experimental traces were processed using custom-written MATLAB scripts. Simulations to extract the magnetic parameters were performed using EasySpin (version 5.2.35)^51^.

The dependence on the applied microwave power of the peak-to-peak EPR signal amplitude A_pp_ measured at the *g*_⊥_ component (Extended Data Fig. 5b) was investigated to characterize the relaxation behaviour of the detected paramagnetic species. Experimental data were modelled according to the equation^52^

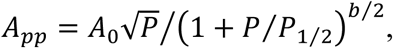

in which *A*_0_ is a constant that depends on the number of spins, *P* is the applied microwave power, *P*_1⁄2_ is the half-saturation power (the microwave power at which the saturation factor *s* = 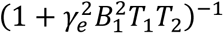 is equal to ½, where *T*_1_ and *T*_2_ are the spin-lattice and spin-spin relaxation times respectively), and *b* describes the homogeneous (*b* = 3: linewidth dominated by electron spin relaxation) *vs.* inhomogeneous character of the EPR line (*b* = 1: linewidth dominated by unresolved hyperfine interactions).

A calibration curve was derived to determine the approximate number of spins present in the EPR sample. Samples of Cu^II^(H_2_O)_6_ in 2 M NaClO_4_ were prepared at known concentrations (50, 100, 150 and 200 µM) and measured in the same EPR quartz tubes (Suprasil, 1.1 mm ID, 1.6 mm OD) used for the cable bacteria samples. For each sample, 10 µL were dispensed into a quartz capillary, resulting in a fill height of approximately 10 mm. The accuracy of the volume was verified by comparing the weight difference between the filled and the empty tubes. As for the cable bacteria samples, the Cu^II^ standards in the Suprasil tubes were loaded inside the wider diameter (2 mm ID, 3 mm OD) CFQ quartz tube and positioned at the centre of the EPR resonator. The spectra were measured at 10 K under strictly non-saturating conditions (8 µW microwave power) using a field modulation of 10 G at 100 kHz, a magnetic field sweep rate of 5.96 G/s, a resolution of 1.02 points/G, a conversion time of 163.84 ms and a time constant of 40.96 ms.

For the determination of the spin numbers in the cable bacteria samples the spectra recorded at the lowest microwave power, namely 1 mW, were used. The double integral of the CW-EPR signal was rescaled by the microwave power, modulation amplitude and measurement temperature according to^53^

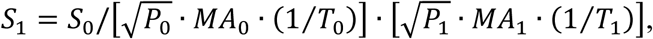

where the subscript 0 refers to the measurement conditions of the cable bacteria samples (*P*_0_ = 1 mW, *MA*_0_ = 20 G, *T*_0_ = 12 K) and the subscript 1 refers to the measurement conditions of the Cu^II^ standards (*P*_1_ = 8 µW, *MA*_1_ = 10 G, *T*_1_ = 10 K). The equation above takes into account that the EPR signal intensity of the chosen standards strictly follows the Curie behaviour, *S* ∝ *T*^-1^.^54^ The number of spins N_spin_ in each sample was deduced from the final calibration curve (Extended Data Fig. 5d, Extended Data Table 2). The height of the cable bacteria pellets was approximately 1 mm whereas the fill height of the Cu^II^ standards was 1.0 cm. As both fill heights are small compared to the full width at half maximum of the resonator mode (∼2.0 cm), the effect of the shape of the resonator profile could be neglected.

To determine the total filament length, the clump of filaments inside each EPR tube was deposited onto a microscope slide within a droplet of mQ. This droplet was dispersed and distributed over multiple (∼10) microscopy slides to disentangle clumps thus enabling the imaging of individual filaments. The microscope slides were dried, and each slide was imaged using a Zeiss Axioplan 2 Imaging microscope with QImaging EXi Blue camera to create a tiled scan of the whole microscopy slide. This scanned images for the different microscopy slides were analysed with ImageJ to attain the total filament length L_fil_ that was present within the EPR tube. The filament length was multiplied with the number of fibres N=68 (Extended Data Fig. 1c), to arrive at the total fibre length L_fibre_ = N_fibre_*L_fil_. The fibre length was multiplied with the Ni content per length of fibre X_Ni_ (as determined by nXRF; Supplementary Table 3) to arrive at the number of Ni atoms N_Ni_ within the EPR tube. Finally, the number of spins in each sample was divided by the number of Ni atoms to arrive at the spin to Nickel ratio N_spin_/N_Ni_ (the calculation procedure is summarized in Extended Data Table 2).

### X-ray Absorption Spectroscopy (XAS) at the Ni K-edge

Square pieces (5 x 5 mm) of polytetrafluorethylene (PTFE) were glued onto a 3D-printed sample holder. Dense clumps of native cable bacteria and fibre skeletons were deposited onto each square, air dried and fixed by polyimide tape (Kapton). Reference compounds included nickel(II)oxide (NiO; TCI Europe N1265), nickel(IV)oxide (NiO_2_; Sigma Aldrich 72262-5G), six monomeric NBDT compounds (same as for Raman spectroscopy; Extended Data Table 1) and two polymeric NBDT compounds (pNi(ett) and p(TTFtt); Extended Data Table 1). All reference compounds were in powder form and diluted with cellulose binder for pellet pressing. Dilution for optimal X-ray absorption to minimize self-absorption was calculated using the XAFSmass software^55^. Calculated amounts were weighed on an electronic precision balance and cellulose was added to a final weight of 60 mg. Powders and binder were mixed by grinding with a pestle and mortar for 15 minutes. Next, the mixtures were pressed into 1 mm thick pellets with a diameter of 8 mm using a pressure of 5 tons. Finally, the pellets were inserted in a metal sample holder and held in place with polyimide tape.

Ni K-edge XAS spectra were recorded at the I20 Scanning beamline of the Diamond Light Source synchrotron^56^, which operates at a ring energy of 3 GeV and current of ∼200 mA. The I20 beamline uses a Si (111) four-bounce monochromator with an energy resolution of 1.5 × 10^−4^ (ΔE/E), resulting in an energy resolution of 1.25 eV at the Ni K-edge (8333 eV), and mirror focusing that provides a beam spot size of 250 μm × 400 μm (height × width). All spectra were recorded in fluorescence mode due to the dilute nature of the samples, and under cryogenic conditions (9 K, pulsed-tube top-loading He cryostat) as to limit beam damage. A 64-element monolithic Ge detector with Xspress4 modular digital pulse processing system measured fluorescence photons from the sample perpendicular to the X-ray beam. The incident beam energy was calibrated to 8,333 eV using a Ni foil. Reference spectra were measured over a range from 8,200 to 8,900 eV in steps of 0.2 eV using an acquisition time of 1 s. Cable bacteria samples were examined in a range from 8,200 to 8,900 eV with steps of 5 eV in the pre-edge region and 0.2 eV in the edge region and acquisition time of 3 s per step. Multiple replicate scans were acquired per sample and the effect of radiation damage was carefully monitored. The acquired XAS spectra were normalized and merged using “Athena” module of the Demeter XAS processing software (version 0.9.26)^57^.

## QUANTIFICATION AND STATISTICAL ANALYSIS

### Image analysis and processing of SEM, TEM and nXRF

Image analysis was performed in Fiji/ImageJ^49^. To derive the ridge compartment width from SEM images of native filaments, line transects were drawn perpendicular to the filament, which revealed consecutive intensity maxima corresponding to ridge crests. The ridge compartment width was measured as the distance between two ridge maxima, and the mean and standard deviation were calculated for a number of n replicates. TEM cross-section images were analysed in ImageJ to determine the fibre diameter within individual ridge compartments. To this end, the fibre core that appeared less stained (Fig. 1d) was delineated via the Freehand Line tool and the surface area A was calculated. The fibre diameter was calculated as 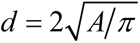. The mean and standard deviation are reported for a number of n replicates.

The elemental areal mass density maps resulting from nXRF mapping were processed within ImageJ and R software. In the elemental mappings of fibre skeletons, we determined (1) the mean ridge width from the Ni map (i.e., the distance between two ridge maxima), and (2) the Ni and S areal mass density of the fibre skeletons, which then provides the S/Ni ratio. In the mappings of fragmented fibre skeletons, we determined (1) the Ni content of individual fibres and (2) the S/Ni ratio of individual fibres. Error propagation was implemented via standard expressions. Mean and standard deviations are reported.

### Density Functional Theory (DFT) simulations

DFT calculations were performed to predict the electronic ground state of different structural models of Ni_3_(ett)_4_ oligomers with different end-capping groups (Extended Data Fig. 6). DFT calculations were performed using the Turbomole code^58^ implementing different density functionals: PBE^59^, OPBE^60^ and B3LYP^61,62^, in combination with Grimme’s D3 dispersion correction^63,64^ including Becke-Johnson dampening, and either the def2-SVP or def2-TZVP basis-set^65^. The geometries of the oligomers were initially optimised as closed-shell singlets before stability analysis^66^ was used to check the stability of the closed-shell Kohn-Sham (KS) wavefunction with respect to open-shell instabilities. For those cases where the closed-shell singlet KS wavefunction was found to be unstable, the geometries of the oligomers were additionally optimised for a set of possible open-shell spin states (open-shell singlet, triplet, quintet).

Additionally, time-dependent DFT (TD-DFT) simulations of the Ni K-edge were performed for pNi(ett) oligomers with three Ni centres (Ni_n_(ett)_n+1_ with n = 3; end capping as in structure III of Extended Data Fig. 6) using the ORCA DFT package^67^. The geometries of the simulated molecular structures were optimized using the B3LYP density functional^61^ in combination with the dispersion correction of Grimme’s method^63,64^ until convergence was achieved within 10^-8^ a.u. The Ahlrichs’ def2-TZVPP basis-set was chosen for the simulations after comparison of the ground-state variational energy against both the Pople’s style and Dunning’s correlation consistent basis sets^65^. The ZORA approximation was used to account for the spin-orbit coupling while the RIJCOSX approximation was included to speed up the calculations. The generalized-gradient PBE functional, hybrid B3LYP, PBE0 functionals, and long-range separated hybrid functional ωB97X-D3 functional^68^ were employed in the TD-DFT calculations within the Tamm-Dancoff approximation formalism for the simulation of XANES spectra. The simulated XANES spectra show the results obtained with the ωB97X-D3 functional, which proved to be most accurate in predicting the energies of the excited states and the relative intensities of XAS features. All simulated spectra have been convoluted with a Gaussian broadening, representing the Ni 1s core-hole lifetime of 1.5 eV.

## Supplementary Information

### 1. Supplementary discussion

#### 1.1 Quality of fibre skeleton extraction

Fibre skeletons attain a flattened, cylindrical shape (Extended Data Fig. 1), which retains the conductive fibres, embedded in parallel on top of a basal carbohydrate-rich sheath^15,23^. The quality of the extraction procedure was verified by Raman microscopy, TEM and AFM imaging. EDTA is known to be a performant chelator of metals, but previous work has shown that the EDTA extraction at 1 mM does not affect the Raman signal intensity of the NiBiD complex, thus suggesting that the structure remains unaltered^15,16^. This is confirmed here. Extracted fibre skeletons display the characteristic Raman fingerprint of the NiBiD complex (Extended Data Fig. 4a), but do not show any marked signal of cytochromes, which are removed during the SDS-EDTA extraction^16^. Likewise, the SDS-EDTA extraction procedure reduces the biomass of native filaments by removing cytoplasmic and periplasmic material (Extended Data Fig. 1c,d). AFM imaging confirmed that the height of the fibre skeletons was substantially reduced compared to native filaments (Extended Data Fig. 1e,f), and fell within the 250-350 nm range previously measured for fibre skeletons^5,15^. The cartwheel structure in the cell-cell interfaces remains largely intact upon extraction, thus producing elevated regions within the AFM images of fibre skeletons (Extended Data Fig. 1f). Moreover, electrical characterization with patterned micro-electrodes (procedure as in Ref. 6) showed that fibre skeletons produced from the JX3-16 clonal culture remained highly conductive after extraction.

#### 1.2 NanoXRF imaging of native filaments and fibre skeletons

Extended Data Fig. 2 compares the nXRF mappings of native filaments with fibre skeletons. The piece of native filament imaged shows four adjacent cells separated by three cell-cell interfaces with slightly enlarged diameters at the cell-cell interfaces (Extended Data Fig. 2 a,c,e,g). The fibre skeleton section also displays four adjacent cells separated by three cell-cell interfaces, but cells are shorter (Extended Data Fig. 2 b,d,f,h). Both native filaments and fibre skeletons show a clear enrichment of Ni in the periplasmic fibres (Extended Data Fig. 2 a,b), thus demonstrating that the fibres account for most of the Ni in the cell and that other Ni-containing enzymes in periplasm and cytoplasm must be less abundant. In comparison, the S mapping displays a different pattern. In the native filament, S appears relatively homogeneously distributed across the filament (Extended Data Fig. 2d). In contrast, the fibre skeletons show a clear enrichment of S in both the periplasmic fibres and the cartwheels (Extended Data Fig. 2c). This suggests that protein-derived sulphur (from cysteine and methionine residues) is prominently present in the S mapping of native filaments, though providing a homogeneous S distribution. In the fibre skeletons, the specific S-signal in fibres and cartwheel could result from either a local accumulation of S-rich amino acids or from the S-containing NiBiD complex. Both native filaments and fibre skeletons show a clear enrichment of Cu in the cell junctions (Extended Data Fig. 2e,f), thus suggesting sizable Cu accumulation in the cartwheels. In contrast, native filaments show a homogeneous distribution of Fe across the cell (Extended Data Fig. 2g), likely originating from cytochromes distributed in cytoplasm and periplasm, while fibre skeletons show an overall low abundance of Fe with some localized hotspots, likely originating from poly-P inclusions (Extended Data Fig. 2h). The Ni, S and Fe mappings obtained here for intact filaments of *Ca*. Electrothrix gigas (Extended Data Fig. 2a,c,e) are consistent with elemental analyses recently performed on intact filaments of a different cable bacterium species, *Ca*. Electronema aureum, as reported in Ref 22. This hence suggests that the Ni/S signal in the fibres is present in all cable bacteria.

#### 1.3 The molar S/Ni ratio of fibre skeletons derived from nXRF maps

We used ImageJ to draw transects (n=10) in the Ni mappings of fibre skeletons perpendicular to the filament direction. In these transects, the peaks in the Ni area density (ρA_Ni_^max^) correspond to the individual periplasmic fibres (Supplementary Fig. 1a). The full width at half maximum of the peaks W_1/2_ = 52 ± 15 nm (n = 128), aligns with the fibre diameter as determined from SEM and TEM imaging. Likewise, the interpeak distance δ_p_ = 147 ± 9 nm (n = 182) provides a measure of the so-called ridge compartment width. This value is slightly larger than the value derived from SEM imaging of native filaments (134 ± 3 nm), and likely originates from the fact that the SDS-EDTA extraction widens the fibre skeleton filaments, and hence slightly increases the interpeak distance.

To arrive at the Ni and S areal mass density across the fibre skeletons (ρA^FS^), we selected large regions of interest (ROI) within the mappings of fibre skeletons (Fig. 1e; Extended Data Fig. 2 b,d), thereby avoiding areas that are non-representative for fibres, e.g., cell junctions where S is not only associated with the fibres but also with the cartwheel, or sections where the fibre skeleton was folded, thus providing a higher S density. To arrive at the background concentration (ρA^BG^), we chose suitable ROIs outside the filament. The mean areal mass densities (ng mm^-2^) of Ni and S were obtained by averaging across all ROIs (n=4).

This analysis provided a mean (background-corrected) area density of Ni (0.067 ± 0.009 ng Ni m^-2^) and S (0.427 ± 0.019 ng S m^-2^) across the fibre skeletons (table S4). In the extraction process that produces the fibre skeletons, cytoplasmic and membrane proteins are removed. Accordingly, we assume that any cytoplasmic Ni-containing enzymes are removed, and so all the Ni in the fibre skeletons belongs to the Ni/S complex embedded in the fibres. This was confirmed by the Raman data (Fig. 2a, Extended Data Fig. 4a) and XAS data (Fig. 2c), which show “clean” Ni/S complex spectra where all modes can be attributed to NiBiD, thus indicating that other Ni-containing enzymes cannot be present in large amounts. As such, we can multiply the calculated mean Ni area density (0.067 ng Ni m^-2^) with the interpeak distance δ_p_ (147 nm), to obtain the amount of Ni that is present per unit length of fibre: 9.9 ± 1.9 x 10^-12^ ng nm^-1^. Accounting for the atomic mass of Ni (58.69 Da) and Avogadro’s number (6.022 x10^23^), this translates into 101 ± 20 nickel centres per nanometre of fibre (Supplementary Table 1). An analogous calculation for S provides the mass of S that is present per unit length of fibre (6.27 ± 0.67 x 10^-11^ ng nm^-1^), which corresponds to a total number of 1177 ± 69 S atoms per unit length of fibre (Supplementary Table 1). The molar S/Ni ratio of a fibre skeleton thus becomes 11.6 ± 3.0 (Supplementary Table 1). It is important to note that this value does not represent the S/Ni ratio of the NiBiD complex, as the sulphur in the fibre skeletons does not exclusively originate from the complex, but also arises from S-containing amino acids (methionine and cysteine) in proteins that make up the fibres as well as other proteins that can be present in the fibre skeleton (see discussion below).

#### 1.4 Ni content of individual fibres derived from nXRF maps

Probe sonication of fibre skeletons enabled nXRF imaging of various fragments, which show colocalization of Ni and S in isolated fibres (Fig. 1g,i; Supplementary Fig. 2) and colocalization of Cu and S in the cartwheels (Fig. 1h,i; Supplementary Fig. 2). We performed image analysis to assess the Ni content of these individual fibres, accounting for two effects: (1) fibres may be flattened onto the surface while drying, thus changing their cylindrical form into an elliptical shape; (2) the X-ray beam focused on the sample has a finite size (43 nm vertical x 33 nm horizontal, full-width at half maximum FWHM), thus resulting in a dispersion effect (point spreading). We adopted a deconvolution approach to compensate for this point spreading effect. In the case of an infinitely small beam, a transect across an elliptical-shaped wire (perpendicular to the wire’s direction) will generate the following theoretical area density profile:

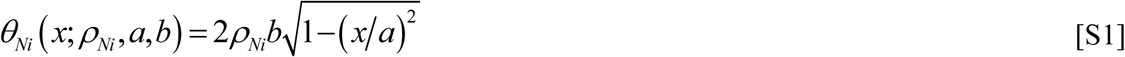

In this, *θ _Ni_* represents the area density of the element at hand, *x* is the distance along the transect, *a* and *b* are the major and minor semi-axes of the fibre’s elliptical cross-section, and *ρ_Ni_* denotes the elemental volumetric density. To simulate the impact of a finite beam size, we implemented a Gaussian point spread function (PSF) with a window parameter w representing the full-width at half maximum (standard deviation of Gauss curve: *σ* = *w*/2.35). The predicted smoothed area density profile is obtained through convolution with the PSF:

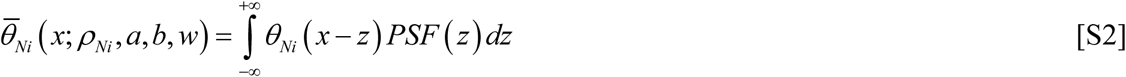

Supplementary Fig. 3b provides an example simulation of the blurring effect on a single fibre. We employed ImageJ to extract the Ni area density for cross-sectional transects across individual fibres (Supplementary Fig. 3a; n=5). For this, we selected fibres that remained isolated on the substrate, i.e. not touching or crossing any other fibres, as to ensure that the recorded area densities truly originated from a single fibre. The function 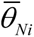 ( *x*) was fitted to these transect data applying the non-linear least squares function *nls* in the R programming software, using *ρ_Ni_*, *a* and *w* as fitting parameters. To reduce the parameter space, we set the minor semi-axis to covary with the major semi-axis, i.e.,. *b* = *r*^2^ *a* with *r* = 25 nm the radius of the original fibre cylinder. This way, we assumed that the fibres keep their original volume when flattening on the surface. Non-linear least squares fitting of 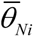 ( *x*; *ρ_Ni_*, *a*, *b*, *w*) (Supplementary Fig. 3c) provided the FWHM beam parameter w = 41 ± 4 nm, which was highly consistent with the known dimensions of the nXRF beam (FWHM = 43 nm). The fitted elliptical shape parameters (a = 38 ± 3 nm; b = 17 ± 2 nm) suggest a flattening of the fibre skeleton upon drying on the substrate.

The best model fit provides a volumetric Ni concentration of *^ρ^_Ni_* = 63 ± 2 mol m^-3^ (Supplementary Table 2). Accounting for the cross-sectional area of a fibre (A = πab =1963 ± 302 nm^2^), we obtained X_Ni_ = 75 ± 15 nickel centres per nanometre of fibre (Supplementary Table 2). This value is slightly lower than the value derived above for fibre skeletons (101 ± 20). This can be explained by the fact that the fibre skeletons form a double layer (cylindrical cell envelope that is flattened, see Extended Data Fig.1b), and so part of the Ni in the lower layer may contribute to the recorded signal in the upper layer. For this reason, we used X_Ni_ = 75 ± 15 # Ni nm^-1^ in all further model calculations as the best estimate of the Ni content of the fibres.

#### 1.5 The molar S/Ni ratio of individual fibres derived from nXRF maps

The nXRF area density maps for S are of poorer quality and more noise-prone compared to the Ni maps, and so the S content of the fibres could not be assessed in the same way as was done above for the Ni content of the fibres (i.e. via direct image analysis of nXRF maps of individual fibres). Therefore, we implemented an alternative procedure, which calculates the S/Ni ratio by means of regression. The nXRF maps of Ni provide a high signal-to-noise ratio and hence a clear outline of the fibres (Supplementary Fig. 3a). Therefore, Ni maps were thresholded in ImageJ to select the image part covered by fibres (the “fibre region”), and this mask was subsequently imposed on the corresponding S map (Supplementary Fig. 4). In both images, Ni and S area densities were extracted on a pixel by pixel basis from the “fibre region”, and background concentrations were subtracted. To arrive at the background concentration (ρA^BG^), suitable ROIs were selected outside the fibre region. The background-corrected areal densities of Ni and S were then analysed by weighted orthogonal regression using the “deming” R package. Supplementary Fig. 4 provides a representative example of the procedure for one image; the procedure was applied to n=7 separate images in total. The slopes provided a mass-based S/Ni ratio of 4.0 ± 0.9, which is equivalent to a molar S/Ni ratio of 7.3 ± 1.6 (Supplementary Table 2). Multiplying this molar S/Ni ratio with the Ni content of the fibres (75 ± 15 nm^-1^), we estimated that the fibres contained X_S_ = 546 ± 229 sulphur centres per nanometre of fibre (Supplementary Table 2). As already mentioned above, one should note that these sulphur centres not only arise from the embedded NiBiD complex, but also from S-containing amino acids (methionine and cysteine) in the proteins that make up the fibres.

#### 1.6 Fibres contain a braided network of electron dense ribbons

HAADF-STEM imaging was performed on cell envelope fragments that were obtained via probe sonication of fibre skeletons. These images reveal electron-dense features that meander along the longitudinal axis of the fibres (Fig. 1k; Supplementary Fig. 5,7). HAADF-STEM selectively collects electrons scattered through very large angles, thus providing increased mass contrast, with regions enriched with heavier elements appearing brighter in the image. The fragment imaged in Supplementary Fig. 5a consisted of parallel fibres (apparent as whiter zones) that were kept together by an underlying sheath (most likely consisting of peptidoglycan). The central zone contains multiple wavy lines that are electron-dense (Supplementary Fig. 5a). These individual ribbons are distinguishable as a series of electron dense dots forming a “pearls on a string” pattern (see magnification in Supplementary Fig. 5b,c). ImageJ analysis indicates that the electron-dense ribbons are ∼1.5 nm in diameter (Supplementary Fig. 5d).

#### 1.7 The molar S/Ni ratio of NiBiD based on HAADF-STEM-EDX

HAADF-STEM-EDX elemental mapping of the cell envelope fragment depicted in Fig. 1k shows a clear enrichment of Ni and S within the centre of the fibres (Supplementary Fig. 6). Line scan profiles of nickel and sulphur show clear coinciding peaks, which match the location of the electron-dense channels (Supplementary Fig. 6c). Line scans also reveal a sequence of consecutive carbon (C) and nitrogen (N) maxima, indicative of the protein that makes up the fibres (Supplementary Fig. 6c). However, the FWHM of the Ni and S peaks is smaller than that of N peak (Supplementary Fig. 6c), thus demonstrating that Ni and S is indeed restricted to the centre of the fibres. Cu and S show an enrichment in larger patches at the fringes of the cell envelope fragment, which are likely remnants of cartwheel structures. Fe does not display a marked fibre pattern (Supplementary Fig. 6a). Orthogonal regression of the Ni and S along the line scan provides a molar S/Ni ratio of 3.0 ± 0.1 (Supplementary Fig. 6d).

HAADF-STEM-EDX imaging at larger magnification shows the electron-dense ribbons within each fibre in greater detail (Supplementary Fig. 7). The associated EDX maps confirm that these electron-dense ribbons are particularly enriched in Ni and S (Supplementary Fig. 7a,e). Line scan profiles of Ni and S show coinciding peaks and a clear colocalization in the centre of the fibres (Supplementary Fig. 7c,g). Orthogonal regression reveals a strong positive linear relationship between Ni and S, providing a molar S/Ni ratio of 2.9 ± 0.5 (Supplementary Fig. 7d,h; Supplementary Table 3; n = 3 replicate images). Assuming that the S signal is almost exclusively derived from the electron-dense ribbons, this value can be interpreted as the molar S/Ni ratio of the embedded NiBiD complex.

#### 1.8 Compositional model of fibre skeletons, fibres and nanoribbons

The above analysis has provided independent values for the molar S/Ni ratio of (1) the entire fibre skeleton (Supplementary Table 1), (2) the individual isolated fibres (Supplementary Table 2), and (3) the nanoribbons embedding the NiBiD complex (Supplementary Table 3). As noted above, these molar S/Ni ratios differ because S is not only contained within the NiBiD complex, but also arises from S-containing amino acids (methionine and cysteine) in proteins that make up the fibres and the rest of the fibre skeleton. To assess the consistency of the obtained molar S/Ni ratios, and to arrive at the best estimate for the S/Ni ratio of the nanoribbons, we constructed a model of that accounts for protein-derived sulphur present within the fibre and the rest of the fibre skeleton (Supplementary Table 3).

As suggested by recent cryoET observations^22^, the fibres contain a network of protein strands that form a matrix in which the electron dense nanoribbon channels are embedded. Adopting a mean protein density of 548 Da/nm^3^ as recorded for bacterial protein fibrils^69^, and a porosity of 0.50 (i.e. half of the fibre volume is made up of protein fibrils), the protein mass per length of fibre (∼50 nm diameter) can be calculated as 5.82 x10^5^ Da nm^-1^, which translates into 5.290 amino acids (AA) nm^-1^, if we adopt a mean molecular weight of 110 Da per AA. Microbial proteins typically contain between 2-4 % of S-based amino acids^69^. If we adopt a value of 3.8% (i.e. on the upper part of the range, thus assuming S-rich peptides), then the fibre protein incorporates 201 ± 21 S atoms per nanometre. In addition, it has been estimated that the fibres themselves only account for 30% of the whole biomass of the fibre skeletons^15^. Accordingly, we estimate that 523 ± 148 S nm^-1^ are present in the additional proteins that make up the fibre skeleton (using the same percentage of S-containing amino acids as above). This way, we estimate that 39% of the S in the fibre skeletons originates from NiBiD containing nanoribbons (454 #S nm^-1^), 17% originates from protein sulphur in the fibres (201 #S nm^-1^) and the remaining 44% (523 #S nm^-1^) originates from sulphur in the additional proteins of the fibre skeleton. The estimated values for the Ni and S present in the NiBiD complex, S in fibre protein and non-fibre protein, and the corresponding S/Ni ratios, are summarized in Supplementary Table 3. In this model, we assume that all the nickel is concentrated in the complex, and so no other Ni-containing proteins are present in the fibre skeleton.

Our model analysis thus provides three independent estimates for the molar S/Ni ratio of the nanoribbons, which are consistent with each other and range between 3.0 and 4.6 (Supplementary Table 3). The value obtained from HAADF-STEM-EDX is lower than the values derived from nXRF mappings, which may signify a bias occurring during HAADF-STEM-EDX, as electron beam irradiation tends to liberate the lighter elements first (i.e., favouring S release relative to Ni), thus lowering the recorded molar S/Ni ratio. Yet overall, the nXRF and HAADF-STEM-EDX results are consistent within the boundaries of uncertainty, thus suggesting that the molar S/Ni ratio of the NiBiD complex within the nanoribbons most likely ranges between 3 and 5.

#### 1.9 Raman microscopy of nickel bis(1,2-dithiolene) complexes

The Raman spectra of fibre skeletons were compared to those of 6 monomeric NBDT complexes (Ni(II)(mnt)_2_, Ni(III)(mnt)_2_, Ni(II)(dmdtb)_2_, Ni(II)(dtb)_2_, Ni(III)(dmit)_2_, and Ni(III)(dttdt)_2_; structures in Extended Data Fig.3b) and two polymeric NBDT compounds (pNi(ett) and pNi(TTFtt); structures in Extended Data Fig.3c). The fibre skeletons and polymeric NBDT compounds were examined at 4 incident laser wavelengths (405 nm, 532 nm, 785 nm, 1064 nm), the monomeric complexes were investigated at three wavelengths (405 nm, 532 nm, 785 nm). In the NIR region, the monomeric NBDT complexes suffered from substantial radiation damage, even at low incident laser power, and as a result, it was impossible to collect spectra with the 1064 nm laser. For the polymeric compounds pNi(ett) and pNi(TTFtt), we noted an effect of ageing (i.e., the time since initial organic synthesis) on the Raman spectra, as intensified peak broadening was observed with increasing age of the sample (as seen in Extended Data Fig.4b for the Raman spectrum of pNi(ett), which was analysed 12 months after synthesis). These ageing effects are attributed to oxidation and accumulating structural disorder of the polymeric strands with time^20^. Accordingly, when comparing spectra. one should be attentive to these ageing effects as well as variation induced by different synthesis and purification procedures. Therefore, we specifically compared the Raman spectra of the NiBiD complex in fibre skeletons to the spectra of freshly synthesized pNi(ett) and pNi(TTFtt) (the spectra for newly synthesized pNi(ett) and pNi(TTFtt) are displayed in Fig. 2a; the Raman spectrum of “aged” pNi(ett) is given for reference in Extended Data Fig.4b; no spectra for “aged” pNi(TTFtt) were recorded)

The Raman spectra of fibre skeletons (Extended Data Fig.4a) shared a strong commonality with those of the monomeric and polymeric reference compounds, thus corroborating the proposition that the Ni-complex has an NBDT-like character. All spectra show a set of common peaks originating from vibrations in the Ni(C_2_S_2_)_2_ core of the NBDT complex (Extended Data Fig.4), which can be assigned to υ(Ni-S) stretching (340 – 400 cm^-1^), and C-S stretching coupled to ring deformation (480 – 530 cm^-1^) (Supplementary Table 5). However, the commonality between the spectra of fibre skeletons and polymeric NBDT compounds is markedly greater than between the spectra of the fibre skeletons and the monomeric NBDT compounds. The latter particularly lack the thiocarbonyl radical (C=S⸱) peaks within the 1150-1200 cm^-1^ region (Extended Data Fig.4). The coordination polymers pNi(ett) and pNi(TTFtt) generate Raman spectra with resonant peaks at near-identical frequencies as the NiBiD complex (Fig. 2a), producing two intense low-frequency peaks near 366 cm^-1^ and 494 cm^-1^ assigned to Ni-S stretching and C-S stretching coupled to ring deformation respectively, which are visible at all 4 wavelengths (Extended Data Fig.4a,b). The relative intensity of these modes also changes in a similar way with the incident laser wavelength: the Ni-S stretching mode (365 cm^-1^) decreases in intensity towards shorter incident wavelengths, while the C-S vibrational mode (495 cm^-1^) shows the opposite behaviour (Supplementary Table 6).

The characteristic Ni-S stretching and C-S vibrations are present in the Raman spectra of monomeric reference compounds Ni, but they occur at different wavenumbers than in NiBiD, and the polymeric pNi(ett) and pNi(TTFtt) (Extended Data Fig.4; annotations in Supplementary Table 5). In general, the Raman spectra of the monomeric NBDT compounds are more complex as they contain additional modes originating from the R substituents (nitrile, benzyl groups) on the 1,2-dithiolene ligands (see molecular structures in Extended Data Fig.3b). The mid-frequency region of the monomeric NBDT spectra contain peaks related to these R substituents, overtones of the low-frequency modes, C=C stretching, and combinations of vibrational modes^24^. In contrast, the spectra of the fibre skeletons, pNi(ett) and pNi(TTFtt) display a limited number of peaks (Fig. 2a, Extended Data Fig.3a,b), thus implying fewer vibrations and a simpler NBDT structure. As such, the Ni/S complex cannot feature nitrile groups or aromatic rings, which would produce intense Raman signals not seen in the spectra^25^.

The high similarity at lower wavenumbers (<500 cm^−1^) between the Raman spectra of NiBiD and those of pNi(ett) and pNi(TTFtt) suggests an intrinsic similarity in structure. The Raman modes observed in the 1,100 – 1,250 cm^-1^ region corroborate this. The spectra of pNi(ett) and pNi(TTFtt) contain two distinct peaks (1,158 cm^-1^, 1,189 cm^-1^; Fig. 2a) attributed to thiocarbonyl radicals (C=S⸱)^20^. These modes are also prominently present in the spectra of the NiBiD complex (1,146 cm^-1^, 1,181 cm^-1^; Fig. 2a), which include an additional peak (1,219 cm^-1^) that may originate from terminal ligations (e.g., cysteine links) not present in pNi(ett) and pNi(TTFtt). These radical modes are prominently present upon excitation with long (785 nm, 1,064 nm) wavelengths (Fig. 2a), but are absent in the green (532 nm) spectra (Extended Data Fig.4a,b), thus suggesting they require absorption in the Near Infrared (NIR) region.

Strong absorption in the Near Infrared (NIR) region is rather unusual in biological molecules^70^, unless they include large, aromatic systems^71,72^. However, the NiBiD complex cannot contain aromatic rings, as aromatic modes (C-C or C=C stretching and ring breathing) would produce characteristic intense Raman signals^25^ that are not seen in the NiBiD spectra. Note that pNi(ett) and pNi(TTFtt) also do not incorporate aromatic rings, but still exhibit strong NIR absorption because of polymerization, which creates an extensive conjugated, planar NBDT system that enables low-energy transitions in the NIR region. The observation of strong NIR resonance (Fig. 2a; Extended Data Fig.4a) therefore suggests that the NiBiD complex is not monomeric, but rather forms a larger complex, in which several Ni centres are interconnected by dithiolene ligands to attain a larger conjugated complex. This is further substantiated by the fact that the thiocarbonyl radical modes do not appear in the 785 nm spectra of the monomeric compounds (Extended Data Fig.4c-f), which lack sufficient conjugation to enable strong NIR absorption.

#### 1.10 Polarized Raman microscopy

To investigate anisotropic Raman scattering, we performed polarized Raman spectroscopy on fibre skeletons oriented in the x-direction using X- and Y-polarized light, while measuring the parallel and cross-polarized components of the Raman scattered light. This way, we collected Raman spectra of the NiBiD complex in four conformations: Z(X,X)Z̅, Z(X,Y)Z̅, Z(Y,Y)Z̅, and Z(Y,X)Z̅ (Extended Data Fig.9c). The signature Ni complex modes (367 cm^-1^, 496 cm^-1^, 1146 cm^-1^, 1181 cm^-1^, 1219 cm^-1^ – see Ref. 16) were most intense when fibre skeletons were positioned horizontally (X) on the substrate and excited by X-polarized light. Analysis of the polarization of the Raman scattered light revealed that the intensity of the parallel component Z(X,X)Z̅ was highest, while the perpendicular component Z(X,Y)Z̅ was markedly lower (Extended Data Fig.9).

Calculation of the depolarization ratio (ρ) (i.e., the ratio of the intensity of perpendicular versus the parallel component of the Raman scattered light) shows that the vibrational modes of the Ni/S complex are highly polarized (ρ < 0.25). The Si-mode of the substrate (520 cm^-1^) showed similar behaviour since the excitation laser polarization was aligned with one of the wafer’s crystal axes. Another mode that showed differences in polarization-dependent scattering was the Amide I mode (1660 cm^-1^). The depolarization ratio of Amide I was 0.25, which corresponds well to reported ratios of ρ = 0.2 - 0.3 in proteins^73^. An overview of the calculated depolarization ratios is given in Supplementary Table 7.

When fibre skeletons positioned in the X-direction were excited with light polarized along the Y-axis, we obtained much lower Raman signals, while the Si-mode (520 cm^-1^) remained unaffected (Extended Data Fig.9b). Analysis of the polarization of Raman signals of the NiBiD complex revealed that the perpendicular component Z(Y,X)Z̅ was more intense than the parallel Z(Y,Y)Z̅ component (Extended Data Fig.9b). This demonstrates that the NiBiD complex scatters more Raman photons along the direction of the conductive fibres irrespective of the polarization. While anisotropic Raman scattering has been observed in biostructures with an important degree of order and directionality, like collagen fibrils^74^ and spider silk^75^, the magnitude of anisotropic Raman scattering as observed here appears exceptional for a biosystem.

#### 1.11 Electron Paramagnetic Resonance spectroscopy

EPR signal recording and spin quantification requires consideration of the temperature window in which the analysis is conducted. At a too high temperature broadening of resonances occurs due to enhanced electron spin relaxation, while at lower temperatures the signal intensity increases because of an increased thermal polarisation of the EPR transition but microwave power saturation may also take place, leading to distortions. As a result of these two mechanisms, there is a temperature region in which the detection of the EPR signal is optimal (i.e., the measurement temperature should be as low as possible with the avoidance of saturating conditions)^76^. To this end, we ensured that our spectra were acquired under non-saturating conditions. Extended Data Fig. 5b demonstrates that the EPR signal is only marginally saturated even at high microwave power (20 mW) and low temperature (12K) and thus results from a relatively fast relaxing species.

As shown in Fig. 2b, the EPR signal is clearly distinguishable (i.e., rises high above the noise level) and can be suitably reproduced by a model of an isolated S = ½ spin system. The paramagnetic character is further supported by the temperature dependence of the recorded EPR signal (Extended Fig. 5e) which follows the Curie-Weiss behaviour S = C/(T-θ), where S is the double integral of the CW-EPR signal and both C (Curie constant) and θ (Weiss temperature) are constants^77^. The very good quality of the linear fit of 1/S against T (Extended Fig. 5e) is consistent with the absence of sizeable electron-electron couplings (exchange interaction and/or zero-field splitting)^78^. Combined with the DFT simulations (see discussion below and Extended Figure 6), these findings are consistent with (weakly coupled) S = ½ spin centres that are mostly localised on the terminal units of the NiBiD complex.

The observed S = ½ EPR signal cannot arise from Ni, because the EPR spectra of Ni^1+^ or Ni^3+^ are characterised by a much larger anisotropy and a much greater deviation of the average g values from g_e_ than observed^79^. As such, Ni must reside in the Ni^2+^ oxidation state, which is either EPR silent (if S = 0) or nearly EPR-invisible at X-band frequency (if S = 1)^76^. This is fully consistent with the DFT simulations of the NiBiD oligomers (see below), which indicate that the Ni centres are +2 oxidation state and carry negligible spin. The electronic configuration of Ni^2+^ is [Ar]4s^0^3d^8^. It can form either high-spin (S = 1) or low-spin (S = 0) complexes, depending on the ligand field strength and coordination. Four-coordinate Ni^2+^ complexes tend to adopt a planar configuration with spin state S = 0, provided it is not forced by extremely bulky ligands into a (pseudo)tetrahedral configuration with spin state S = 1. Our Raman and XANES data indicate a square planar geometry, and therefore, S = 0 is the most likely spin ground state. However, to more confidently constrain the Ni spin ground state, one requires investigation at much higher magnetic fields, if observable at all given the large zero-field splitting^80^.

The recorded g = 2.014 value suggests that the radical should be located on an atom heavier than C, N or O, as otherwise it would show a smaller g value^81^. The deviation of g from the free-electron g-value g_e_ can be described by g = g_e_ ± nλ/ΔE, where n is a quantum mechanical coefficient, λ is the spin-orbital coupling constant and ΔE is the energy gap between ground an excited state. Because λ increases for heavier atoms, the deviation from g_e_ is larger for an S-centred radical compared to an O-, C-or N-centred radical (assuming the same or similar ΔE). Such an S-centred radical^82^ is consistent with the DFT calculations, which predict that the spins are localised on the terminal S-containing ligands on either side of the complex (Extended Data Fig. 6e). Likewise, it is consistent with thiocarbonyl radical (C=S⸱) peaks seen in the Raman spectrum.

The radical S = ½ signal persists in spectra recorded at low temperature (12 K) and high microwave powers (20 mW), thus marking an anomalously fast relaxation behaviour for an organic radical^83^ (Extended Data Fig. 5a). Our experimental findings suggest that it’s unlikely that the detected signal originates from an ‘isolated’ radical as it would be heavily saturated under the chosen experimental conditions. The fast relaxation suggests the presence of a nearby spin that mediates the observed fast relaxation^83^. This could be caused by paramagnetic relaxation from a neighbouring S = 1 Ni^2+^ centre, but as argued above, square planar Ni^2+^ complexes typically adopt a low-spin S = 0 state with paired electrons, and not a high-spin S = 1 state. An alternative option is that the observed spin relaxation originates from spins that are located on neighbouring strands within the metal-organic framework.

To examine the possible origin of the radical S = ½ signal further, we performed DFT simulations of NiBiD complexes with three Ni centres, examining different terminal ligands and charge states (Extended Data Fig. 6). These calculations predict that there are two unpaired electrons residing on the terminal ligands, thus providing a complex that forms a biradical (Extended Data Fig. 6e). When the unpaired electrons are far apart, as is the case for NiBiD complexes with many Ni centres (see Extended Data Fig. 6e), the spin–spin interactions become negligible. If the two radical centres are identical (e.g., if the terminal ends of the NiBiD complexes are similar and the molecule is symmetric), the EPR signal will resemble a single radical S = ½, as observed here (Fig. 2b). Our DFT simulations hence indicate that for certain configurations of Ni_n_(ett)_n+1_ (structures III and IV in Extended Data Fig. 6), the electronic structure is consistent with the radical S = ½ signal observed by EPR.

#### 1.12 X-Ray Absorption Spectroscopy

Nickel K-edge measurements of the X-ray absorption near-edge structure (XANES) and Extended X-ray Absorption Fine Structure (EXAFS) were performed to gain more detailed insights into the oxidation state of the Ni centre and the nature of the coordinating ligands in the NiBiD complex. The XAS spectra of native filaments were compared to those of 6 monomeric NBDT complexes (listed in Extended Data Table 1; Extended Data Fig. 3b), two polymeric NBDT compounds (pNi(ett) and pNi(TTFtt); Extended Data Fig. 3c) and the Ni pincer containing enzyme LarA^29^.

Ni K-edge spectra for native filaments were highly consistent between two biological replicates of the same synchroton XAS session, as well as between synchroton XAS sessions conducted at different times, thus demonstrating good reproducibility. While the monomeric NBDT complexes showed good stability over time, the XAS spectra of the polymeric compounds pNi(ett) and pNi(TTFtt) showed an effect of ageing, with marked changes in the XANES spectra since the initial time of synthesis (as also noted for the Raman spectra). To avoid this ageing effect, we compared the XAS spectra of native cable bacterium filaments to the spectra of freshly synthesized pNi(ett) and pNi(TTFtt), as described in Ref. 20 (spectra in Fig. 2a).

The Ni K-edge of NiBiD lies at a different energy than the inorganic Ni reference compounds (Extended Data Fig.7a), but closely resembles that of the monomeric and polymeric NBDT complexes (Fig. 2c; Extended Data Fig.7b). The difference in edge positions is explained by the higher covalency of the S-containing ligands in NBDT complexes, compared to the O-ligands of the Ni oxide reference compounds. Higher covalency contributes to charge neutralization, resulting in a lower effective charge (Z_eff_) on Ni, and hence a shift to a lower rising-edge energy. High covalency also delocalizes the Ni 4p orbitals, which decreases the transition intensity of the main edge. Together, these effects broaden the rising edge in NBDT complexes and shift it towards lower energies^84,85^. Since the Ni K-edge of NiBiD displays the same downward shift and broadening as the NBDT reference complexes, its Ni centre must reside in a similar oxidation state and must be coordinated by similar S-containing ligands.

The Ni K-edge spectrum of native cable bacteria filaments closely resembles that of the monomeric and polymeric NBDT reference compounds. Two peaks can be observed near the Ni K-edge (Fig. 2c; Extended Data Fig.7b). The pre-edge peak at about 8,333 eV can be assigned to a 1s → 3d electron transition, while the shoulder peak, which occurs at ∼8,337 eV, is linked to a 1s → 4p_z_ transition^24,86^. The latter is a typical XANES feature for square planar complexes^24^. In octahedral Ni(II) complexes, both the 1s → 3d and 1s → 4p_z_ transitions are forbidden, while in square-planar complexes they are allowed. As a result, in octahedral complexes the shoulder peak at 8,337 eV is absent, while the pre-edge feature shows a much smaller intensity than in square-planar complexes (orbital mixing by lower symmetry gives some probability for the 1s → 3d transition).

Among the six monomeric NBDT reference compounds investigated, the XAS spectrum from Ni(dmit)_2_ most closely resembles the spectrum of NiBiD (Extended Data Fig. 7c). The spectrum is nearly identical in EXAFS region, and within the XANES region, Ni(dmit)_2_ shows a similar pre-edge peak at about 8,333 eV, and similar characteristic features at ∼8,350 eV and ∼8,363 eV. Still, the Ni(dmit)_2_ spectrum reveals a more pronounced rising shoulder peak at ∼8337 eV and it also incorporates an additional feature at ∼8,345 eV that is not present in NiBiD (Extended Data. Fig 7c). In terms of features, the XAS spectra of pNi(ett) and pNi(TTFtt) are highly similar to NiBiD: they provide a nearly identical EXAFS spectrum (Fig. 2d), while the XANES region (Fig. 2c) shows a weak pre-edge peak at about 8333 eV and similar peaks at 8,350 eV and ∼8,363 eV. The feature seen at ∼8,345 eV in Ni(dmit)_2_ is also not present (Fig. 2c). Notably, the peak at 8,350 eV slightly differs in intensity between NiBiD, pNi(ett) and pNi(TTFtt). However, we observed that this peak showed variation between different samples of cable bacteria, thus suggesting it could be sensitive to O_2_ exposure during sample preparation, and therefore, the observed difference at 8,350 eV does not impose strong structural constraints on whether NiBiD is more similar to either pNi(ett) or pNi(TTFtt). Another marked difference between the XAS spectra of NiBiD and those of the polymeric NBDT is the intensity of the rising-edge shoulder (8,338 eV), which is absent in pNi(TTFtt) and only weakly expressed in pNi(ett). To examine the factors contributing to the formation of this of the rising-edge shoulder, we performed time-dependent DFT simulations of XANES spectra for Ni_3_(ett)_4_-oligomers with different end-capping R groups. Interestingly, both the DFT simulation of Ni_3_(ett)_4_ terminated with 4 thiols (Structure III in Extended Data Fig. 6) as well as Ni_3_(ett)_4_ terminated with 4 cysteine units (Structure IV in Extended Data Fig. 6) display a similar prominent pre-edge feature as seen in the experimental XANES spectra of NiBiD (Extended Data Fig. 7d). This suggests that the rising edge-shoulder is not only present in monomeric NBTD, but also oligomeric Ni_n_(ett)n+1 complexes. However, the intensity of this shoulder decreases when more Ni(ett) units were added (n > 3) in the TD-DFT simulations (data not shown), which could indicate that the polymeric NBTD still consist of longer polymer strands with more Ni centres than NiBiD^14^.

#### 1.13 DFT calculations

DFT calculations with the hybrid B3LYP density functional predict that the electronic ground state of structural models I and II (Extended Data Fig.6a,b) are closed-shell singlets while those of structural models III and IV (Extended Data Fig.6c,d) are open-shell singlets, where the open-shell singlet state lies 0.2-0.3 eV lower in energy than the closed-shell singlet state and the open-shell singlet and open-shell triplet are split by ∼25 meV (Supplementary Table 8). Calculations with both the small def2-SVP and large def2-TZVP basis-sets give very similar results. Löwden spin population analysis of the open-shell singlet and triplet states shows that the majority of the spin in the open-shell states is localized on the sulphur and carbon atoms of the terminal ligands, in line with the character of the highest occupied orbitals shown for structural model IV (Extended Data Fig.6e). The DFT B3LYP calculations for structural models III and IV agree with the experimental EPR results in terms of the open-shell nature of the electronic ground state and the fact that the spins localize on the organic ligands rather than the nickel atoms. DFT calculations using the GGA PBE and OPBE functionals predict that the electronic ground state of all structural models I-IV is closed-shell, which is at odds with the EPR results.

## 2. Supplementary tables and figures

### 2.1 Supplementary tables

**Supplementary Table 1.** Molar S/Ni ratio and associated parameters derived from ImageJ analysis of nXRF elemental mappings of fibre skeletons. See Supplementary Text for the calculation procedure.

| Fibre skeletons | Units | Value |
| --- | --- | --- |
| Ni area density | ng mm<sup>-2</sup> | 0.067 ± 0.009 |
| S area density | ng mm<sup>-2</sup> | 0.427 ± 0.019 |
| Interpeak distance | nm | 147 ± 9 |
| # Ni per fibre length | #Ni nm<sup>-1</sup> | 101 ± 20 |
| # S per fibre length | #S nm<sup>-1</sup> | 1177 ± 69 |
| # molar S/Ni ratio | / | 11.6 ± 3.0 |

**Supplementary Table 2.** Molar S/Ni ratio and associated parameters obtained from ImageJ analysis of nXRF elemental mappings of individual fibres. See Supplementary Text for the calculation procedure.

| Individual fibres | Units | Value |
| --- | --- | --- |
| Ni density | mol m<sup>-3</sup> | 63 ± 2 |
| Ni concentration per fibre length | ng Ni nm<sup>-1</sup> | 7.3 ± 1.4 |
| # Ni per fibre length | #Ni nm<sup>-1</sup> | 75 ± 15 |
| # molar S/Ni ratio | / | 7.3 ± 1.6 |
| # S per fibre length | #S nm<sup>-1</sup> | 546 ± 229 |

**Supplementary Table 3.** Three independent estimates of the molar S/Ni ratio of the Ni/S-complex in cable bacteria. (*) indicate model estimates of how much protein-derived sulphur is present within the fibre and the rest of the fibre skeleton. All other values are directly based on nXRF or HAADF-STEM-EDX data (see text). <u>Model 1</u>: values determined from nXRF mappings of fibres skeletons (fig. S2B,D), sequentially accounting for non-fibre protein S and fibre protein S. <u>Model 2</u>: values determined from nXRF mappings of individual fibres (fig. S4-S6), accounting for fibre protein S. <u>Model 3</u>: estimate of molar S/Ni ratio in NiBiD complex as determined by HAADF-STEM-EDX mappings of nanoribbons in individual fibres (fig. S8-S9). All Ni and S content values are expressed as atoms per nanometre of fibre. S/Ni ratios represent molar ratios.

|  | Model 1<br>nXRF<br>fibre skeletons | Model 2<br>nXRF<br>individual fibres | Model 3<br>HAADF-STEM-EDX<br>individual fibres |
| --- | --- | --- | --- |
| NiBiD complex Ni | 101 ± 20 | 75 ± 15 | - |
| NiBiD complex S | 454 ± 163 | 345 ± 230 | - |
| Fibre protein S | 201 ± 21 (*) | 201 ± 21 (*) | - |
| Total S in fibre | 655 ± 303 | 546 ± 229 | - |
| Non-fibre protein S | 523 ± 148 (*) | - | - |
| Total S in fibre skeleton | 1177 ± 69 | - | - |
| S/Ni ratio Fibre skeleton | 11.6 ± 3.0 | - | - |
| S/Ni ratio Fibre | 6.5 ± 2.5 | 7.3 ± 1.6 | - |
| **S/Ni ratio NiBiD** | **4.5 ± 2.5** | **4.6 ± 4.0** | **3.0 ± 0.5** |

**Supplementary Table 4.** Optimized Raman spectroscopy acquisition parameters for the nickel bis(dithiolene) complexes.

| Laser | Grating (gr/mm) | Power (mW) | Acquisition time (s) | Range (cm<sup>-1</sup>) |
| --- | --- | --- | --- | --- |
| 405 nm | 2400 | ≈ 1.5 | 60 | 100 – 2000 |
| 532 nm | 1800 | ≈ 0.25 | 60 | 100 – 2000 |
| 785 nm | 1200 | ≈ 0.05 | 300 | 100 – 2000 |
| 1064 nm | 830 | ≈ 0.5 | 480 | 100 – 2000 |

**Supplementary Table 5.** Annotation of the principal vibrational modes in the NiBiD and pNi(ett). The source column indicates references used for annotation.

| Annotation | NiBiD | pNi(ett) | Source |
| --- | --- | --- | --- |
| S-Ni-S deformation | 180 cm<sup>-1</sup> | 186 cm<sup>-1</sup> | (16,24) |
| $\nu(\text{Ni-S})_{\text{sym}}$ – Ni-S stretching | 367 cm<sup>-1</sup> | 366 cm<sup>-1</sup> | (16,24) |
| $\nu(\text{C-S})$ + ring deformation | 496 cm<sup>-1</sup> | 494 cm<sup>-1</sup> | (16,24) |
| $\nu(\text{C=S}\cdot)$ – thiocarbonyl radical stretching | 1146 cm<sup>-1</sup> | 1158 cm<sup>-1</sup> | (20,86) |
| $\nu(\text{C=S}\cdot)$ – thiocarbonyl radical stretching | 1181 cm<sup>-1</sup> | 1189 cm<sup>-1</sup> | (20, 86) |
| $\nu(\text{C=S}\cdot)$ – thiocarbonyl radical stretching | 1219 cm<sup>-1</sup> | / | (20) |

**Supplementary Table 6.** The ratio between the Raman peaks at 367 and 496 cm^-1^ for NiBiD and 366 and 494 cm^-1^ for pNi(ett).

| Laser | NiBiD complex | pNi(ett) |
| --- | --- | --- |
| 1064 nm | 1.45 ± 0.24 | 2.29 ± 0.20 |
| 785 nm | 1.66 ± 0.59 | 2.10 ± 0.14 |
| 532 nm | 0.42 ± 0.05 | 0.54 ± 0.05 |
| 405 nm | 0.40 ± 0.04 | 0.45 ± 0.04 |

**Supplementary Table 7.**
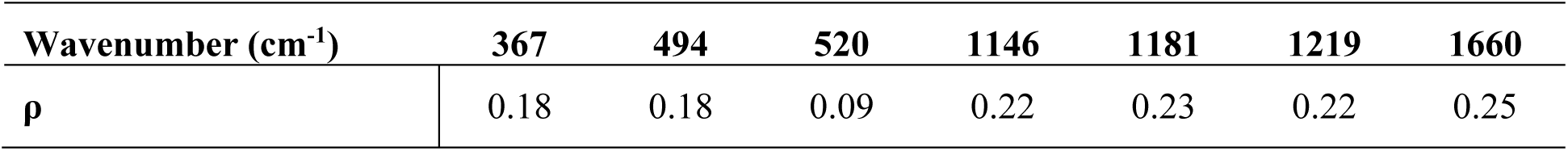
Depolarization ratios (ρ) of key vibrational modes in the Raman spectra of fibre skeletons. Depolarization ratios were calculated as the intensity of the perpendicular versus the parallel component of the Raman signal for each mode. The NiBiD complex modes (367 cm^-1^, 496 cm^-1^, 1,146 cm^-1^, 1,181 cm^-1^, 1,219 cm^-1^) have a low depolarization ratio (< 0.25), indicating they constitute totally symmetric modes. Other modes in the table include Amide I (1,660 cm^-1^) and the Si-mode of the substrate (520 cm^-1^).

| Wavenumber (cm<sup>-1</sup>) | 367 | 494 | 520 | 1146 | 1181 | 1219 | 1660 |
| --- | --- | --- | --- | --- | --- | --- | --- |
| $\rho$ | 0.18 | 0.18 | 0.09 | 0.22 | 0.23 | 0.22 | 0.25 |

**Supplementary Table 8.** Electronic ground state as predicted by DFT modelling for the different structural models defined in Extended Data Figure 6. The ground state is indicated. For those models where the ground state is predicted to be open-shell, additional quantities are calculated: (1) the stabilisation energy (E_stab_) of the open-shell ground state with respect to a structure optimised as a closed-shell singlet, (2) the vertical singlet-triplet gap (E_ST,v_), which represents the splitting in energy between the open-shell singlet and triplet at the optimised open-shell singlet geometry, and (3) the adiabatic singlet-triplet gap (E_ST,a_), i.e., the difference in energy between the optimised open-shell singlet and triplet structures. Calculations are performed with the B3LYP functional and the def2-SVP basis-set (results obtained with the def2-TZVP basis-set are highly similar and placed between parentheses). All values in eV.

| Structure | Ground State | $E_{\text{stab}}$ | $E_{\text{ST,v}}$ | $E_{\text{ST,a}}$ |
| --- | --- | --- | --- | --- |
| I | Closed-shell<br>Singlet |  |  |  |
| II | Closed-Shell<br>Singlet |  |  |  |
| III | Open-Shell<br>Singlet | -0.30 (-0.31) | 0.025 (0.024) | 0.022 (0.022) |
| IV | Open-Shell<br>Singlet | -0.27 | 0.028 | 0.027 |

### 2.2 Supplementary figures

**Supplementary Figure 1.**
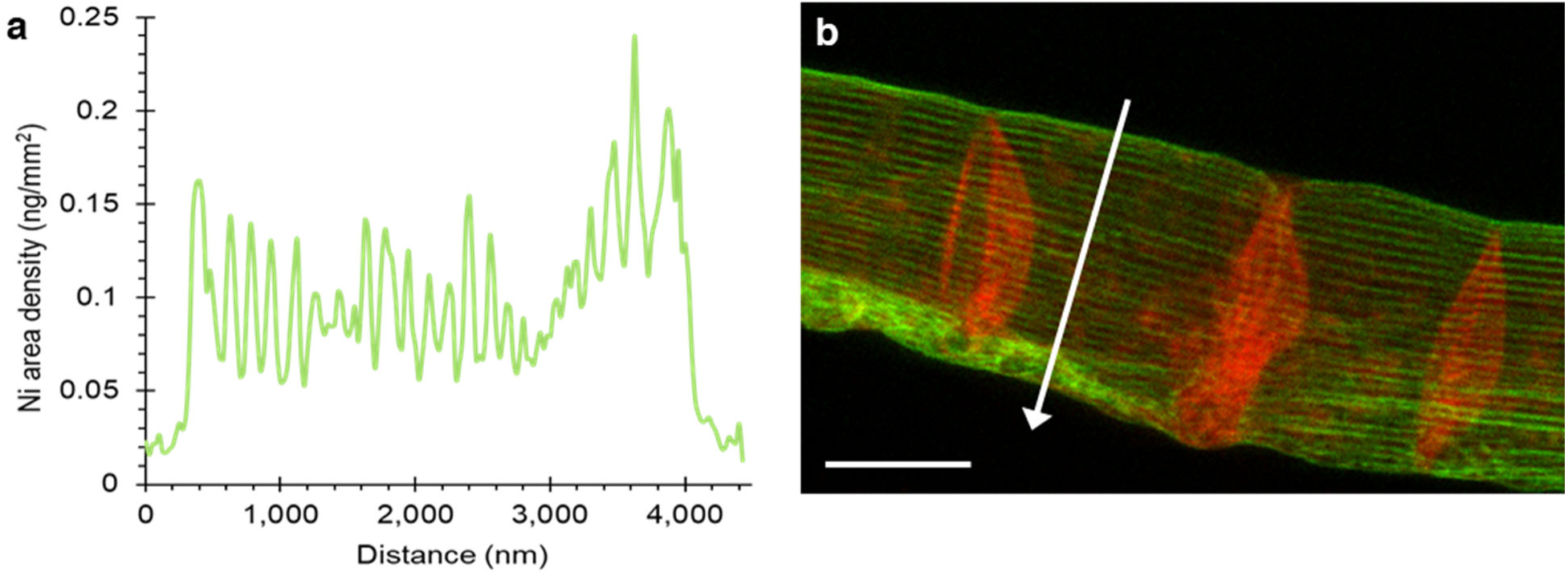
Image analysis of the Ni area map obtained by synchrotron nXRF of a fibre skeleton. **a**, Example of a profile plot of the Ni area density (ρANi) along a cross-section through a fibre skeleton filament (as indicated by the white arrow in panel b). The peaks in the profile correspond to individual periplasmic fibres. **b**, Super position of the Cu (red) and Ni (green) area density of the fibre skeleton. The corresponding maps for individual elements are shown in Fig. 1e and Fig. 1f respectively. Scale bar: 2 µm.

**Supplementary Figure 2.**
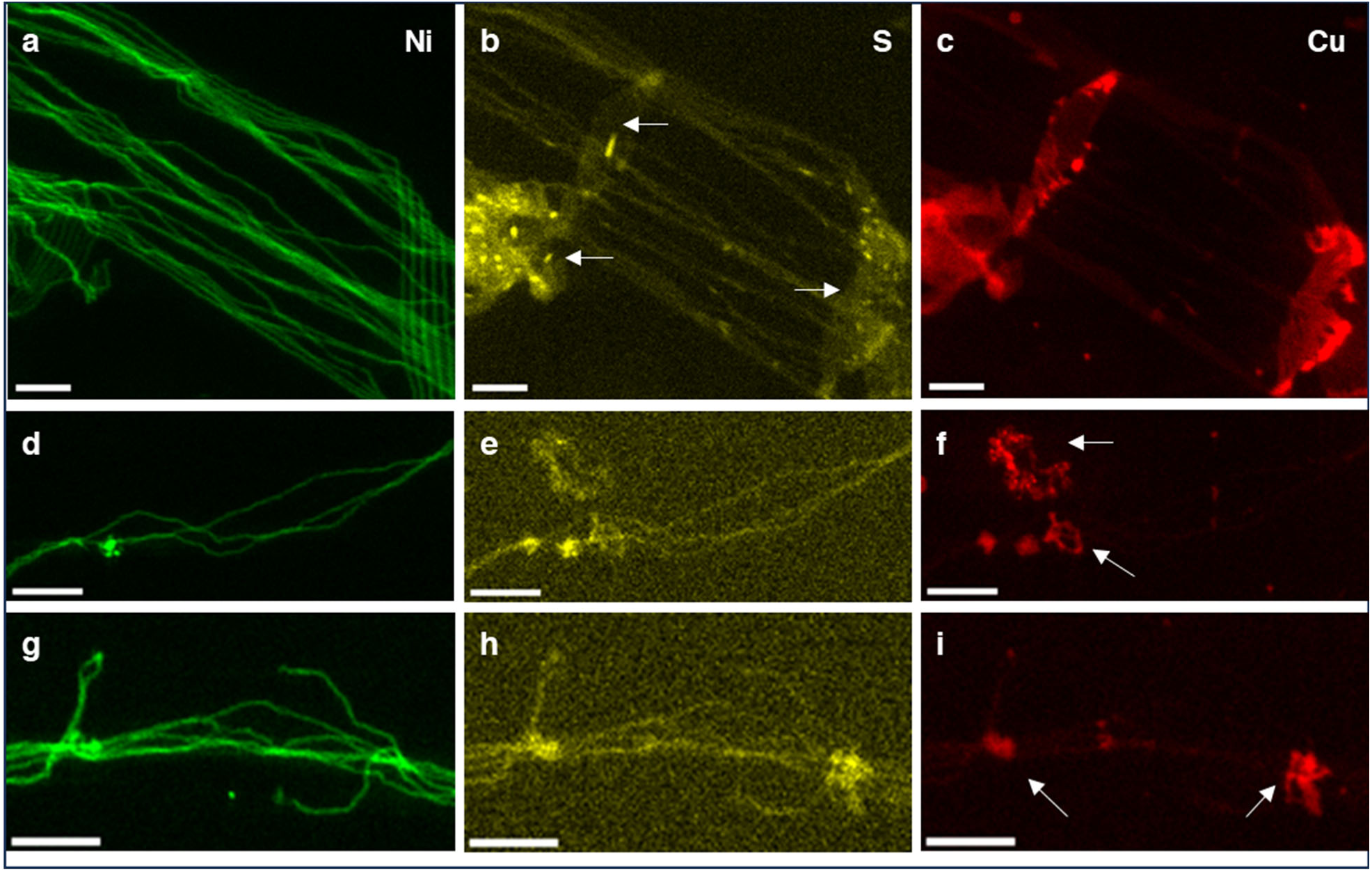
Synchrotron nXRF elemental maps of sonicated fibres skeletons. **a-c**, Area density maps of a fragmented fibre skeleton. Fibre structures are visible in the Ni and S maps. Cartwheel structures are visible in the S and Cu maps (white arrows). **d-f**, Area density maps of three individual fibres, showing colocalization of Ni and S. White arrows indicate a piece of cartwheel debris. **g-i**,. Area density maps of a bundle of individual fibres. White arrows indicate pieces of cartwheel that make a connection between the fibres. All scale bars: 1 µm.

**Supplementary Figure 3.**
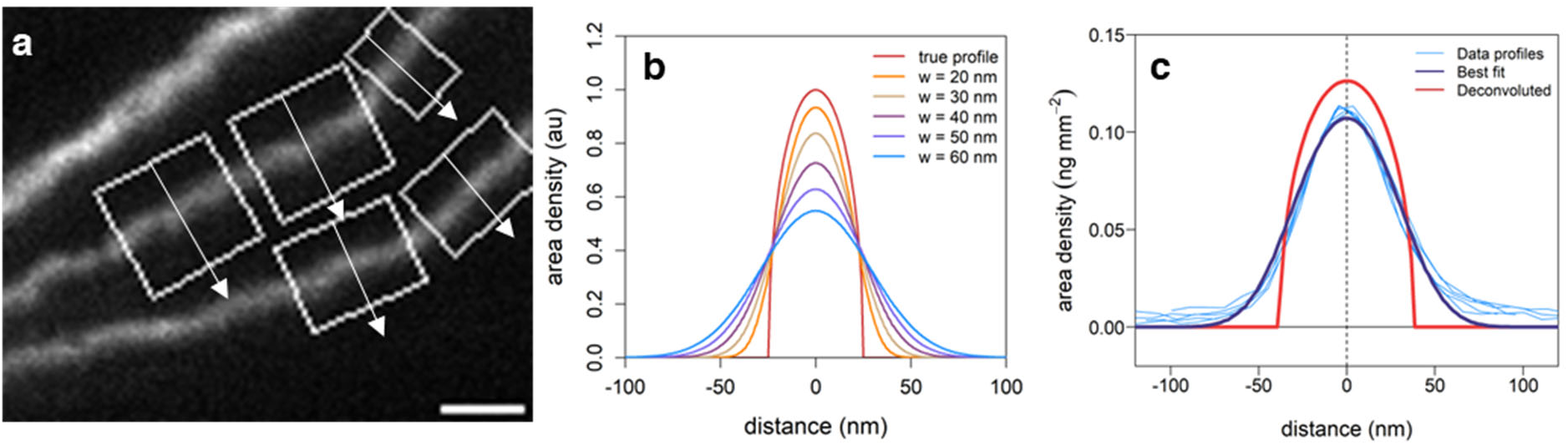
Quantification procedure for the Ni content of individual fibres. **a**, nXRF area density map for Ni depicting three fibres that were loosened from a fibre skeleton by sonication. The rectangular ROIs indicate the location where unidirectional line scans were performed (corresponding data profiles given in panel c). Scale bar: 200 nm. **b**, Model analysis of point spreading on the nXRF mappings due to a finite beam size. The true Ni density profile for a cylindrical fibre (a = 25 nm, b = 25 nm) as calculated by Eq. (1) is given in red. The other profiles simulate the impact of point spreading with a finite beam size. Curves calculated via Eq. (2) using a Gaussian point spread function (PSF) with a window parameter w representing the full-width at half maximum. When w increases, the nXRF mapping will record a more “smoothened” Ni profile. **c**, The measured Ni profiles along the unidirectional line scans in panel a (light blue) are depicted together with best fit by Eq. (2) (dark blue), and its subsequent deconvolution (red).

**Supplementary Figure 4.**
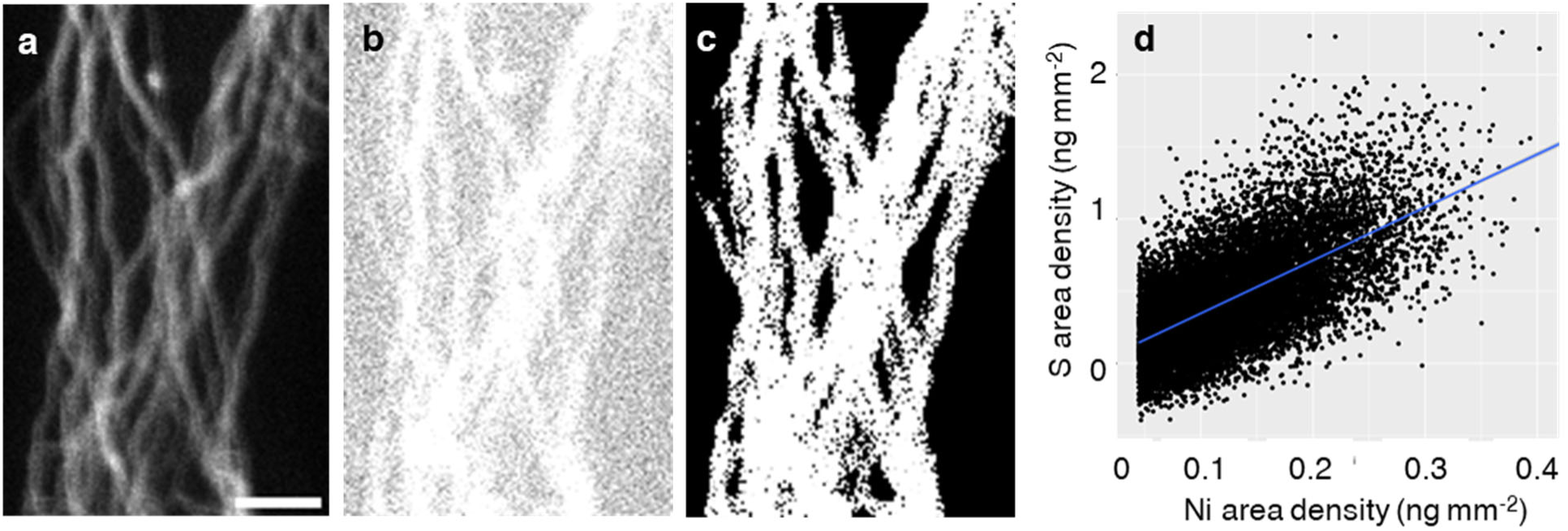
Quantification procedure for the S/Ni ratio of individual fibres. **a**, nXRF area density map for Ni depicting a set of loosened fibres obtained by sonication of a fibre skeleton. Scale bar: 1 µm. **b**, Corresponding nXRF area density map for S. **c**, Thresholded Ni map. White area designates pixels belonging to the fibres. **d**, Scatterplot of the Ni area density versus the S area density for all pixels belonging to the thresholded region in panel c. The blue line denotes the orthogonal regression line from which the molar S/Ni ratio is directly calculated. The images a-d provides a representative example for the calculation procedure of the Ni/S ratio (total of n=7 images were analysed).

**Supplementary Figure 5.**
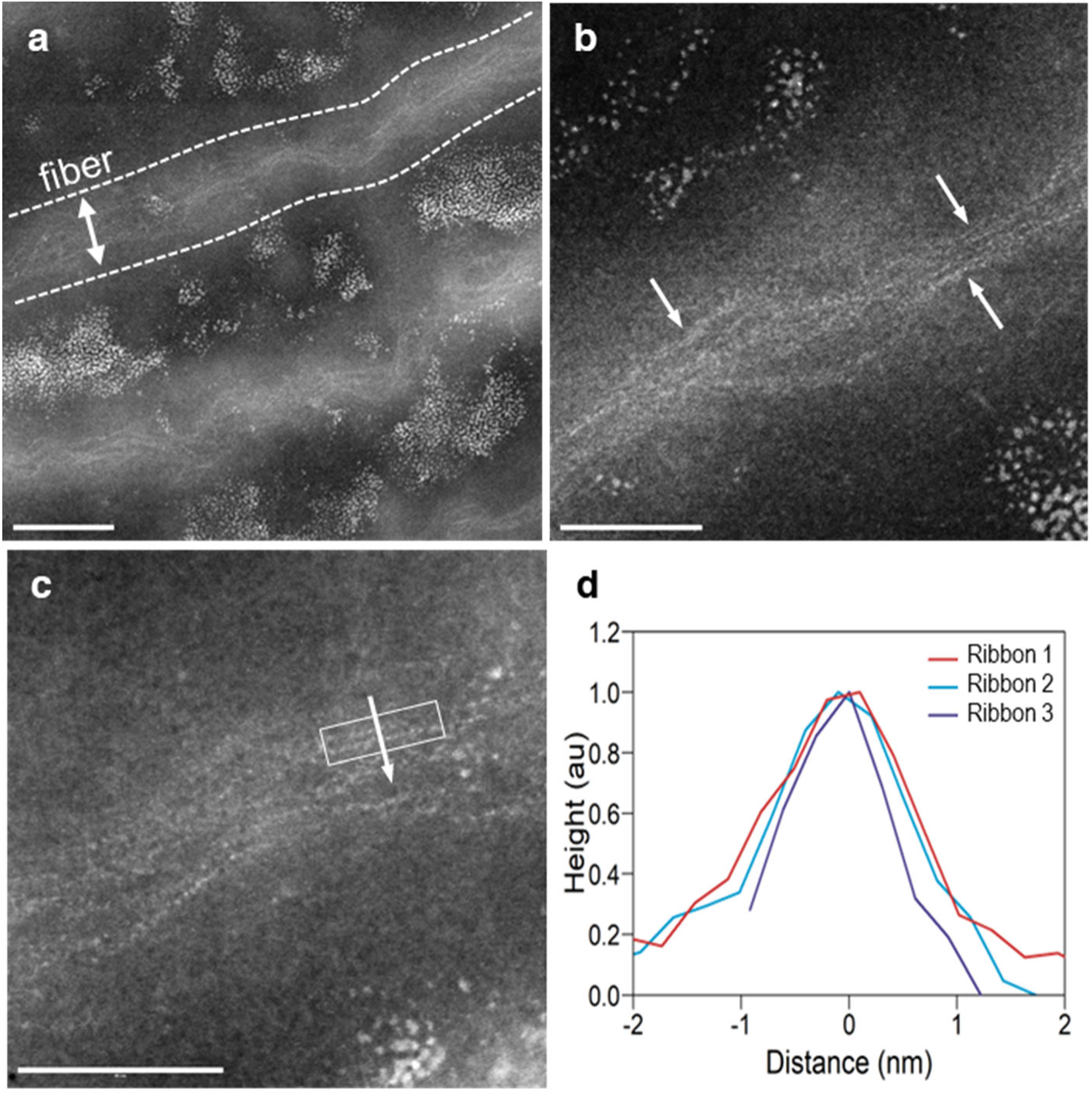
High resolution HAADF-STEM images of fibre skeleton fragments showing individual fibres and nanoribbons. **a**, Cell envelope fragment showing 2 parallel fibres. Electron dense ribbons are apparent as electron-dense lines in the centre of the fibres. Scale bar: 100 nm. **b**, Zoom of single fibre. Individual nanoribbons are distinguishable as a series of electron dense dots forming “pearls on a string”. Scale bar: 50 nm. **c**, Cross-section transection through a single nanoribbon. Scale bar: 50 nm. **d**, Transversal intensity profiles across 3 separate nanoribbons (one example is indicated in panel c). Intensity profiles are centred and rescaled to the maximum intensity.

**Supplementary Figure 6.**
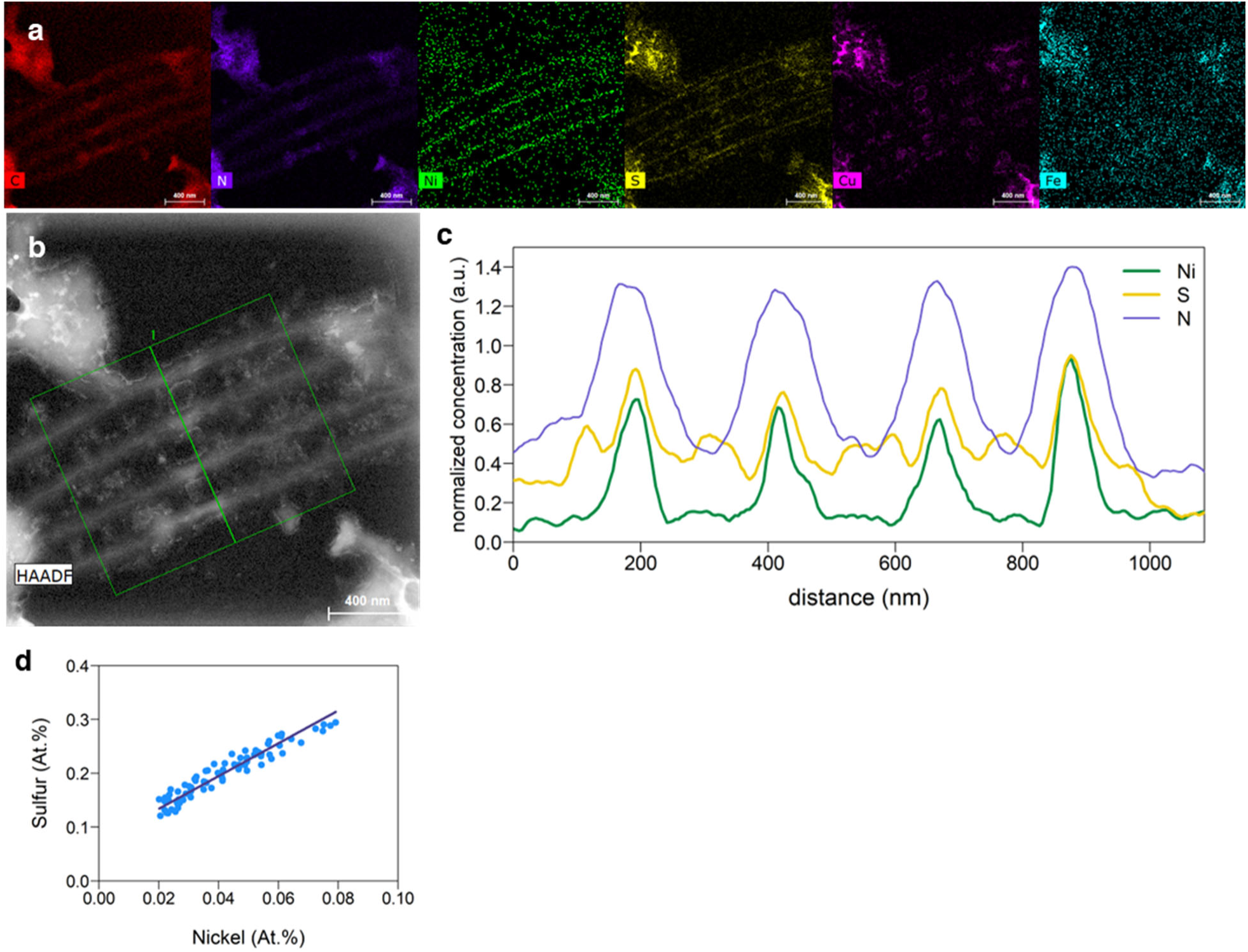
HAADF-STEM-EDX elemental mapping of a cell envelope fragment showing 4 parallel fibres. The fragment was obtained via probe sonication of fibre skeletons. **a**, Elemental mappings obtained by EDX. Different colour panels indicate different elements as indicated. All maps on same scale. **b**, HAADF-STEM intensity image showing the ROI used for line scanning. Scale bar: 400 nm. **c**, Line scan profile of nickel, sulphur and nitrogen revealing the colocalization of the Ni and S in the centre of the fibres (the Full Width at Half Maximum of the Ni and S peaks coincides, but is smaller than that of the N peak). Profiles are normalized for clarity (a.u. = arbitrary units). **d**, Orthogonal regression of the Ni and S along the directional line scan shown in panel b.

**Supplementary Figure 7.**
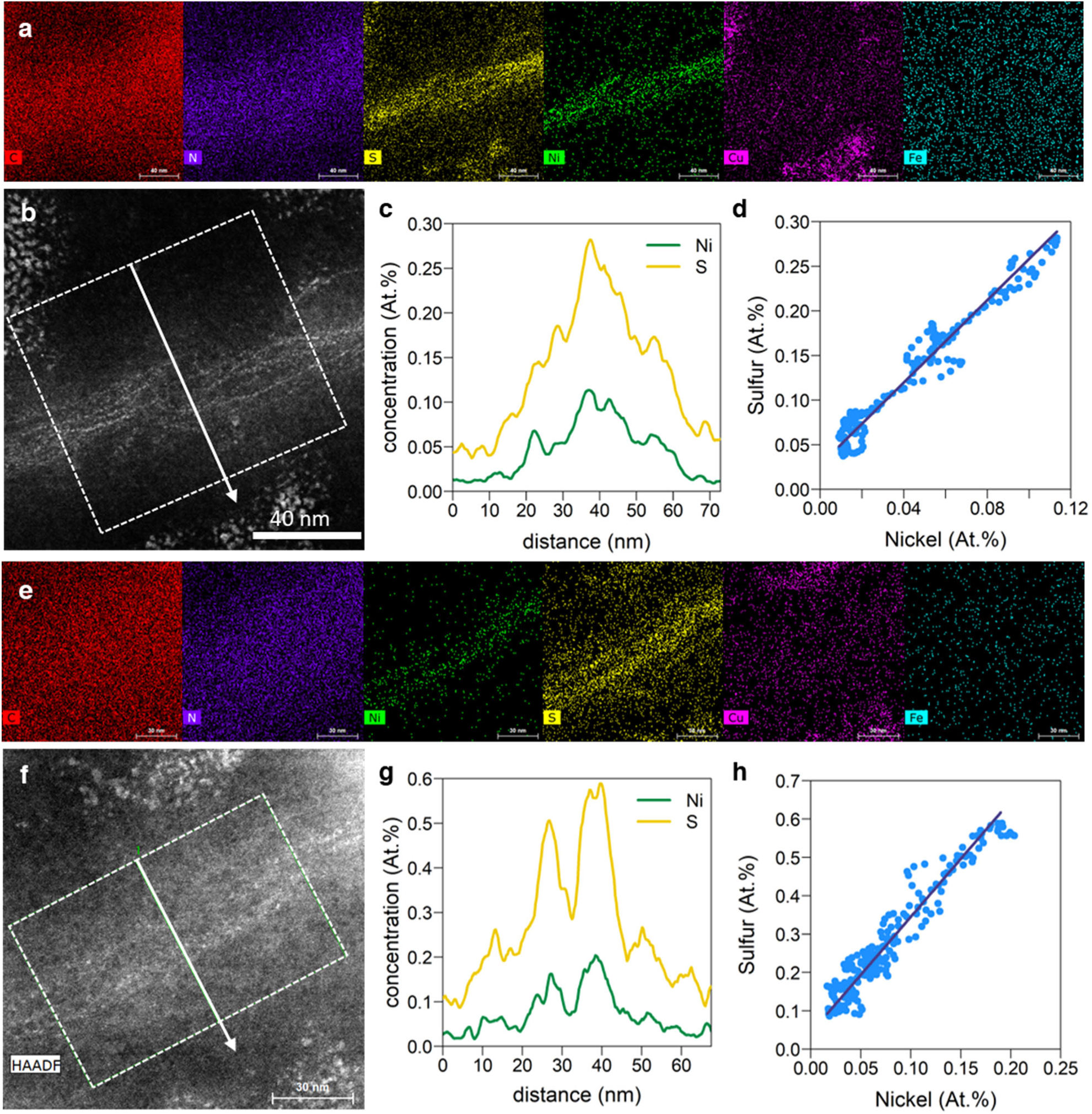
HAADF-STEM-EDX elemental mapping of individual fibres. Two representative examples are shown. **a, e**, Elemental mappings obtained by EDX. Different colour panels indicate different elements as indicated. All corresponding maps are on same scale. **b, f**, HAADF-STEM intensity image showing the ROI used for line scanning. **c, g**, Line scan profile of nickel and sulphur revealing the colocalization of Ni and S in the centre of the fibres. **d, h**, Orthogonal regression of the Ni and S along the line scan shown in panel B and F. The slope of the regression provides the S/Ni ratio of the nanoribbons embedded in the fibre.

